# Conformational landscape and clustering of human type 2 IP_3_ receptor in lipid nanodiscs

**DOI:** 10.64898/2026.01.29.702656

**Authors:** Caifeng Liu, Yu-Jing Lan, Qingyu Tang, Erkan Karakas

## Abstract

Inositol 1,4,5-trisphosphate (IP_3_) receptors (IP_3_Rs) are tetrameric ER Ca^2+^ channels that shape intracellular Ca^2+^ signaling in response to IP_3_, regulating many physiological processes. The structural basis for subtype-specific regulation of three subtypes (IP_3_R-1-3) remains incompletely understood, due to the lack of IP_3_R-2 structures. Here, we determined cryo-electron microscopy (cryo-EM) structures of human IP_3_R-2 in distinct conformations with and without IP_3_, Ca^2+^, and ATP. These structures define the conformational ensembles adopted by IP_3_R-2 and delineate ligand-binding interactions. Comparison with rat IP_3_R-1 and human IP_3_R-3 highlights shared architectural features and isoform-specific differences that underlie subtype-specific functional properties. We also resolved structures of IP_3_R-2 clusters, providing insight into mechanisms of ligand-dependent clustering. Together, these findings establish a structural framework for human IP_3_R-2, linking ligand recognition to conformational transitions and interchannel interfaces, and illuminate how subtype-specific features and clustering may shape cellular Ca^2+^ signaling.

## INTRODUCTION

Calcium (Ca^2+^) is a ubiquitous second messenger that regulates diverse physiological processes, including apoptosis, gene expression, and muscle contraction^1,2^. Ca^2+^ homeostasis is essential for cell viability, whereas its dysregulation can disrupt cellular functions and lead to disease^3–8^. Inositol 1,4,5-trisphosphate (IP_3_) receptors (IP_3_Rs) integrate diverse inputs to shape Ca^2+^ signals by tightly controlling their spatial and temporal characteristics within cells, thereby directing downstream responses^3–6^. Primarily located on the endoplasmic reticulum (ER) membrane, IP_3_Rs are activated upon IP_3_ binding, generated via phospholipase C-mediated hydrolysis of phosphatidylinositol 4,5-bisphosphate (PIP_2_) downstream of G protein-coupled receptors (GPCRs) or receptor tyrosine kinases (RTKs)^3–6^. The resulting channel opening releases Ca^2+^ from ER stores into the cytosol. IP_3_Rs also require Ca^2+^ binding to cytosolic activating sites at nanomolar Ca^2+^ concentrations, whereas higher cytosolic Ca^2+^ concentrations inhibit gating, producing a biphasic dependence of channel activity on Ca^2+ 9–13^. IP_3_R activity is further regulated by small molecules such as ATP^14,15^, posttranslational modifications such as phosphorylation^16^, and interactions with regulatory proteins^17^.

In mammals, the IP_3_R family has three subtypes (IP_3_R-1, IP_3_R-2, and IP_3_R-3), encoded by three different genes (*ITPR1*, *ITPR2*, and *ITPR3*, respectively)^18–20^. All three subtypes share 60-70% amino acid sequence identity and assemble as homo- or heterotetramers, forming one of the largest intracellular ion channel complexes (∼1.2 MDa). Despite this high sequence identity, the subtypes exhibit distinct spatiotemporal expression patterns and physiological properties, resulting in isoform-specific channel behavior. In addition to expression differences, the subtypes differ in key physiological properties: IP_3_R-2 exhibits the highest affinity for IP_3_, whereas IP_3_R-3 has the lowest^3,19,21–23^; IP_3_R-1 and IP_3_R-2 are more sensitive to potentiation by ATP than IP_3_R-3^23–25^. Under comparable conditions, channels composed of IP_3_R-2 tend to support cell-wide Ca^2+^ oscillations, whereas IP_3_R-3-containing channels more often produce large, sustained Ca^2+^ transients with a reduced propensity to oscillate^26–28^.

In recent years, advances in cryo-electron microscopy (cryo-EM) have yielded high-resolution structures of rat IP_3_R-1 (rIP_3_R-1)^29–32^ and human IP_3_R-3 (hIP_3_R-3)^33–36^ in multiple ligand-bound states, illuminating their gating mechanisms. However, structural information for IP_3_R-2 has been notably absent, a gap often attributed to difficulties in expression and purification^23^. To address this, we synthesized a codon-optimized gene for human IP_3_R-2 (hIP_3_R-2) and recombinantly expressed the protein. Following purification and reconstitution into lipid nanodiscs, we determined cryo-EM structures of hIP_3_R-2 in distinct conformations with and without IP_3_, Ca^2+^, and ATP. These structures define the conformational ensembles adopted by hIP_3_R-2, delineate ligand-binding interactions, and reveal consequences for channel gating.

Comparison with rIP_3_R-1 and hIP_3_R-3 highlights shared architectural features and isoform-specific differences that underlie subtype-specific functional properties. We also resolved the structure of hIP_3_R-2 dimer, providing insight into the mechanisms of ligand-dependent (IP_3_ and Ca^2+^) clustering, a phenomenon associated with the formation of Ca^2+^ puffs —defined as localized, transient ER Ca^2+^ release events arising from the coordinated opening of IP_3_Rs^28,37^.

## RESULTS

### Expression, purification, and structure determination of hIP_3_R-2 in lipid nanodiscs

To enable structural characterization of hIP_3_R-2, we synthesized the full-length gene, expressed the protein using Sf9 insect cells, and reconstituted it into MSP1E3D1/DOPC nanodiscs (Fig. 1a). Channel activity was confirmed by planar DOPC/DOPE bilayer recordings. In the presence of IP_3_ and ATP, hIP_3_R-2 exhibited robust single-channel openings with a biphasic dependence on free Ca^2+^ and maximal open probability (Po) at ∼100-200 nM (Fig. 1b-c).

**Figure 1:**
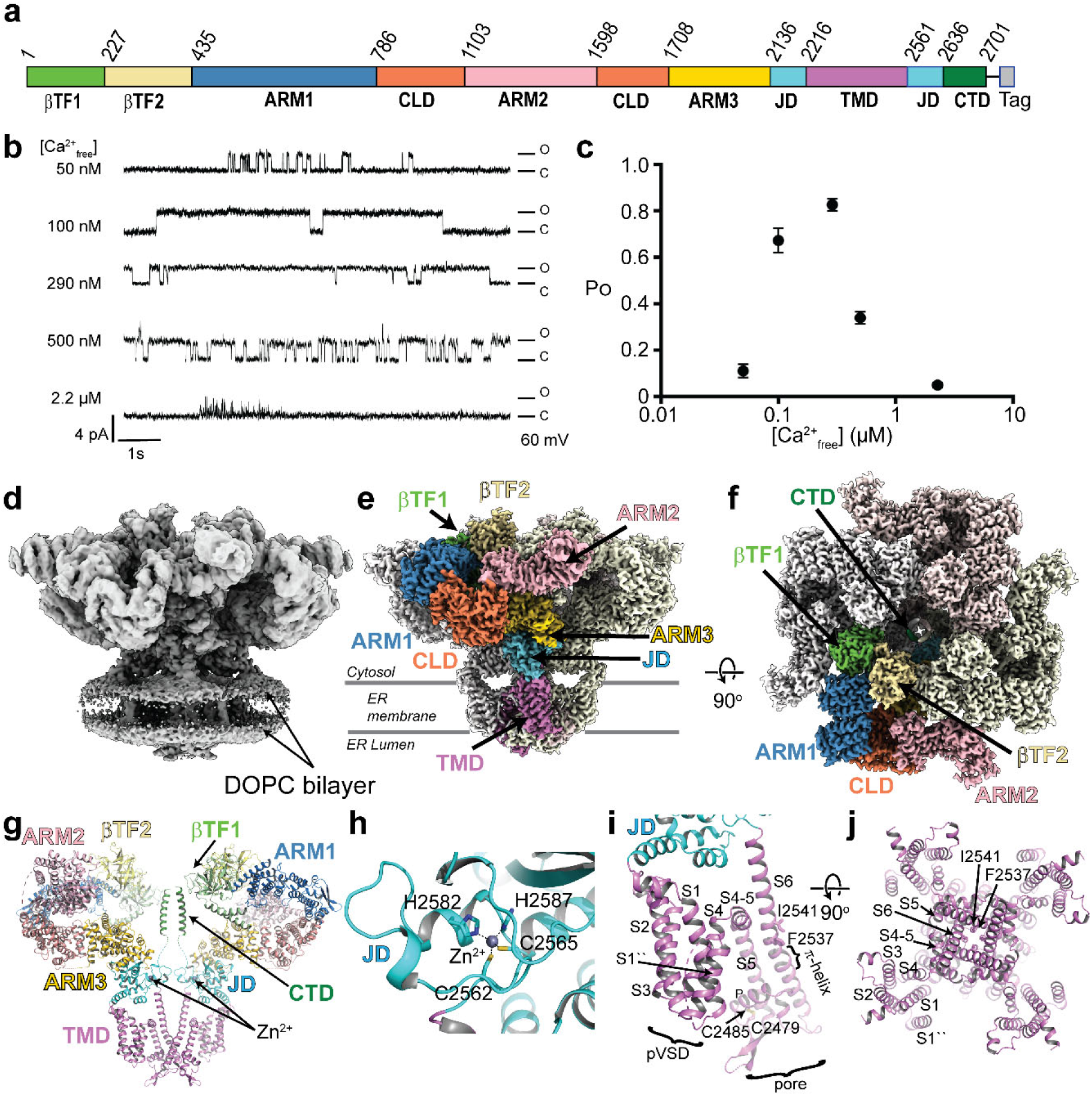
Functional and structural characterization of hIP_3_R-2. **a,** Domain boundaries of hIP_3_R-2 construct. **b,** Representative single-channel traces of hIP_3_R-2 at different Ca^2+^ concentrations. **c,** Plots of open probability as a function of free Ca^2+^ concentrations. Error bars correspond to s.e.m (n ≥6; each independent recording corresponds to a newly incorporated channel). **d,** Unsharpened consensus map of apo hIP_3_R-2. **e-f,** Composite maps of hIP_3_R-2. Each domain in one of the subunits is colored as in panel a. **g,** Ribbon diagram of the apo hIP_3_R-2 showing only two opposing subunits. **h,** The close-up view of the zinc-finger motif. **i,** Close-up view of the TMD (only one subunit is shown). **j**, Close-up view of the TMD as viewed through the pore.

We prepared three cryo-EM samples to capture distinct conformational states under different ligand conditions: (1) ligand-free (apo), (2) with 50 µM IP_3_ and 1 mM ATP (IP_3_/ATP), and (3) with 50 µM IP_3_, 1 mM ATP, and 4 mM CaCl_2_ (IP_3_/ATP/Ca^2+^). Because the protein buffer contained 5 mM EDTA, the calculated free Ca^2+^ concentration for the IP_3_/ATP/Ca^2+^ condition was ∼10 nM (MaxChelator), and the other two conditions were predicted to have negligible free Ca^2+^.

### Structure of apo hIP_3_R-2

Under apo conditions, we identified two major conformational classes, hereafter termed compact and loose, based on the arrangement of the cytosolic domains (Supplementary Fig. 1). The compact class shows well-resolved cytosolic domains forming intersubunit contacts and closely resembles hIP_3_R-3 conformations observed at low Ca^2+ 33–36^. By contrast, the loose class displays weakened or lost cytosolic intersubunit contacts and increased flexibility, resembling the hIP_3_R-3 conformations observed at high Ca^2+ 34–36^. 3D refinement of the particles in the compact class with C4 symmetry yielded a high-quality map at 2.95 Å overall resolution (Fig. 1d and Supplementary Fig. 1). Symmetry expansion and focused local refinements further improved map quality in select regions (Fig. 1e-f and Supplementary Fig. 2). Given the close similarity to published rIP_3_R-1 and hIP_3_R-3 apo structures^31–33,35,36^ and the absence of density for IP_3_, ATP, and Ca^2+^ at expected sites, we designate this reconstruction as the hIP_3_R-2 apo state (Fig. 2a). The loose class, refined without symmetry enforcement, yielded a 4.10 Å map (Supplementary Fig. 1). While its overall architecture is similar to high-Ca^2+^ hIP_3_R-3 conformations^34–36^, this map does not permit confident Ca^2+^ assignment. Refinement strategies that improved the compact class did not markedly improve this class, consistent with increased conformational heterogeneity. We therefore refer to this ensemble as inactive-like and do not infer inhibition.

**Figure 2:**
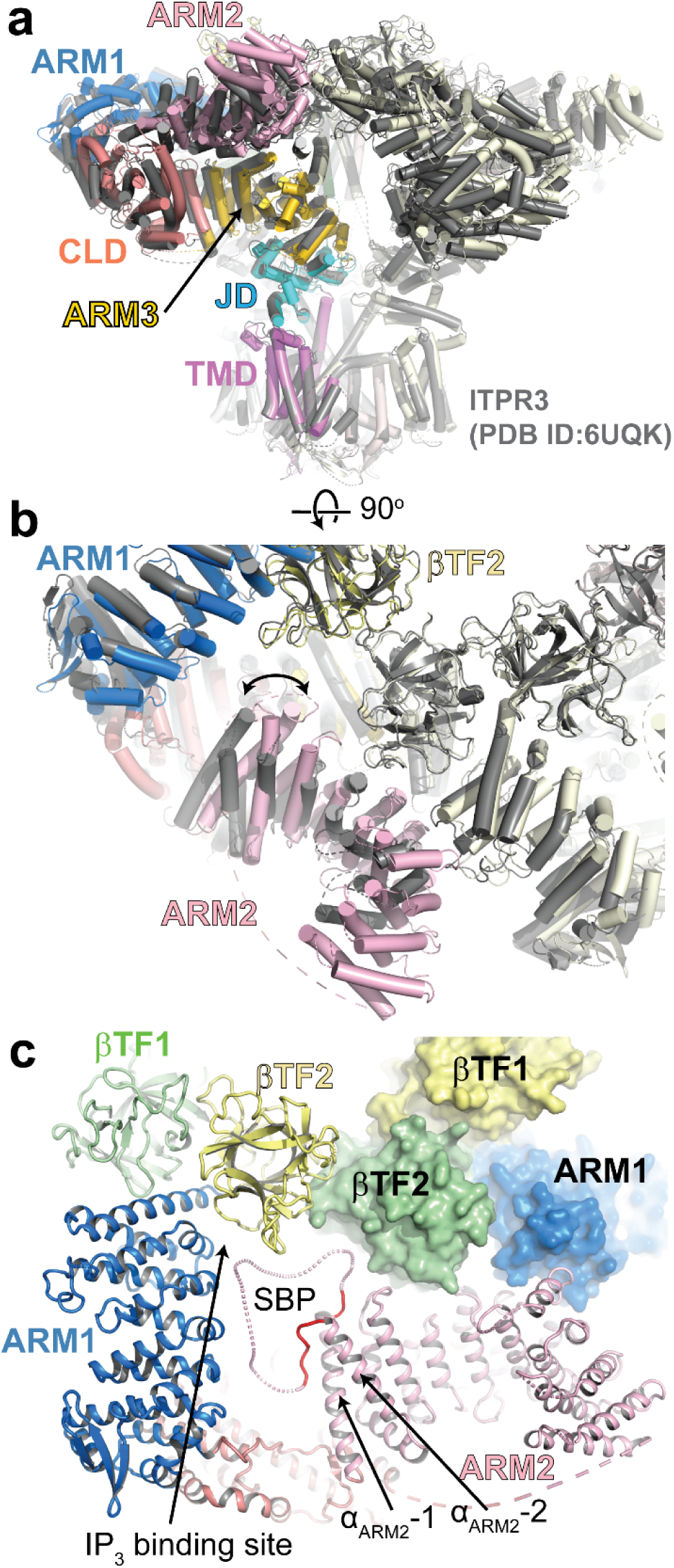
Comparison of the hIP_3_R-2 and hIP_3_R-3 structures. **a-b,** Structures of apo hIP_3_R-2 and hIP_3_R-3 (PDB ID: 6UQK) overlayed on their TMDs. Domains in one hIP_3_R-2 subunit are colored as in Fig. 1, and hIP_3_R-3 structure is colored in grey. b, The same overlay as in panel a, but focusing on the ARM2 domains. **c,** The close-up view of the cytosolic domains of two apo hIP_3_R-2 subunits; one shown in ribbon and another in surface representation. The region that is resolved in hIP_3_R-2 compared to hIP_3_R-3 is colored red. The SBP peptide is not modeled but shown as dashes to highlight potential conformations based on 3D classification results.

The overall architecture of apo hIP_3_R-2 closely matches that of rIP_3_R-1 and hIP_3_R-3, forming a characteristic mushroom-shaped assembly with a cytosolic cap and membrane-embedded stem that constitutes the ion conduction pathway^31–33,35,36^ (Figs. 1d-g and 2a). A small portion of the stem is exposed to the ER lumen and includes glycosylation sites located within flexible, unresolved regions. The channel is a homotetramer with fourfold symmetry around the pore axis (Fig. 1f). Each subunit adopts the same domain organization as in rIP_3_R-1 and hIP_3_R-3, comprising two β-trefoil domains (βTF1 and βTF2), three Armadillo repeat domains (ARM1, 2, and 3), a central linker domain (CLD), a juxtamembrane domain (JD), a transmembrane domain (TMD), and a C-terminal domain (CTD) (Fig. 1e-g). The map quality enabled the modeling of most residues; however, regions with no interpretable density were excluded from the final model. Although the presence of a left-handed coiled-coil helical bundle within the CTD is evident in the cryo-EM map, unambiguous assignment of the residues forming this bundle was not possible. Therefore, we modeled this segment as a poly-alanine.

There are many parallels between hIP_3_R-2 and rIP_3_R-1/hIP_3_R-3. Individual domains superpose on their counterparts with Cα RMSD <1 Å, indicating a conserved fold. As in rIP_3_R-1 and hIP_3_R-3, a zinc-finger motif within the JD coordinates Zn^2+^ via Cys2562, Cys2585, His2582, and His2587 (Fig. 1h). The TMD adopts a fold similar to voltage-gated ion channels, with S1-S4 forming a pseudo-voltage sensor-like domain (pVSD) and S5-S6 forming the pore, connected by the S4-S5 linker helix (Fig. 1i-j). Unlike canonical voltage-gated channels, the cryo-EM map reveals two additional TM helices between S1 and S2. One of these helices (S1″) is well ordered with clear side chain density, whereas the other (S1′) appears flexible and could not be modeled with confidence (Fig. 1i-j). Analogous helices were observed in rIP_3_R-1 and hIP_3_R-3 structures, suggesting that they are a conserved feature of IP_3_Rs^29–31,33–36^. The narrowest region of the pore is formed by Phe2537 and Ile2541 on the S6 (Fig. 1j). In the apo state, the solvent-accessible pore radius at this constriction is about 0.7 Å (calculated using HOLE^38^), consistent with a closed gate. As in rIP_3_R-1 and hIP_3_R-3, the S6 helix contains a π-helical segment on the luminal side of these pore-blocking residues (Fig. 1i).

Despite the high overall similarity, apparent differences emerge in tetrameric assemblies when apo hIP_3_R-2 and apo hIP_3_R-3 (PDB 6UQK^33^) are aligned on their TMDs (Fig. 2a). Most differences are subtle, reflecting rigid-body misalignments among cytosolic domains and likely falling within the intrinsic conformational variability of these regions. The most noticeable difference is in the arrangement of ARM2, which is more tilted toward the adjacent subunit in hIP_3_R-2 (Fig. 2b).

### Self-binding peptide (SBP) dynamics in hIP_3_R-2

We previously showed that a flexible loop within the ARM2 domain of hIP_3_R-3 can occupy the IP_3_-binding site of the same subunit^33^. This loop, termed the self-binding peptide (SBP), competes with IP_3_ binding and likely contributes to the regulation of IP_3_R activation. In hIP_3_R-3, the cryo-EM map revealed substantial density within the IP_3_ pocket; however, its connection to ARM2 was not visible, and a complete SBP model could not be constructed^33^. In the hIP_3_R-2 map, we do not observe a similar density attributable to the SBP at or near the IP_3_-binding site, and the segment linking ARM2-α1 and ARM2-α2 was also unresolved, suggesting high flexibility. By contrast, additional portions of the SBP extending from both ARM2-α1 and ARM2-α2 are visible in hIP_3_R-2: three residues from ARM2-α1 project toward βTF2 of the adjacent subunit, consistent with a potential intersubunit contact, and five residues at the N-terminal end of ARM2-α2 bend by ∼180° and run roughly parallel to ARM2-α2 while forming extensive intradomain contacts (Fig. 2c). The loop between these ordered segments remains disordered.

To better assess SBP occupancy and configurations that may be obscured by averaging in the global maps, we performed 3D classification using a mask encompassing the IP_3_ pocket (Supplementary Fig. 3). Although the resulting maps were insufficient to build the SBP residues unambiguously, they indicate multiple SBP conformations: in approximately one-third of particles, density consistent with the SBP occupies the IP_3_ site (Supplementary Fig. 3a-d); in another third, part of the SBP approaches βTF1 of the adjacent subunit, consistent with a potential intersubunit contact (Supplementary Fig. 3d-g); and in the remaining particles, the SBP is not resolved, consistent with higher flexibility and/or heterogeneity (Supplementary Fig. 3h-j). Together, these observations support a model in which the SBP samples multiple conformations in hIP_3_R-2, including self-occlusion of the IP_3_ pocket and intersubunit engagement, potentially tuning ligand access and allosteric coupling.

### Structure of hIP_3_R-2 in complex with IP_3_ and ATP

As in the apo condition, the IP_3_/ATP dataset yielded compact and loose classes (Supplementary Fig. 4). Sequential 3D classification of compact particles resolved six states: two high-resolution reconstructions (3.31 and 3.33 Å) with apparent fourfold symmetry corresponding to ligand-bound resting and preactivated, and four intermediates (intermediate states 1-4) at 6.22, 6.91, 4.47, and 4.17 Å (Supplementary Figs. 4-5). Intermediate states 1 and 2 are the lowest-resolution maps and were not analyzed further; intermediate states 3 and 4 supported rigid-body fitting of individual domains and capture asymmetric domain arrangements between the resting and preactivated ensembles. The loose class adopts an inactive-like conformation. In all reconstructions, IP_3_ and ATP show well-defined density at their expected binding sites.

In the ligand-bound resting state, the overall architecture closely resembles the ligand-free apo structure despite strong density for IP_3_ and ATP (Fig. 3a). IP_3_ occupies the cleft between the βTF2 and ARM1 domains and induces a modest clamshell-like motion, with ARM1 rotating ∼4° toward βTF2 (Fig. 3b). The most noticeable conformational change is on the loop that harbors Arg269, which moves closer to the IP_3_ molecule and forms a salt bridge with the P5 phosphate of IP_3_. Additional interactions are predominantly electrostatic contacts between basic residues and the phosphate groups of IP_3_: Arg265 (βTF2) with P4; Lys507, Arg510, and Lys569 (ARM1) with P5; and Arg503 and Arg568 (ARM1) with P1. Thr267 (βTF2) and Tyr567 (ARM1) are within hydrogen-bonding distance to P4 and P5, respectively.

**Figure 3:**
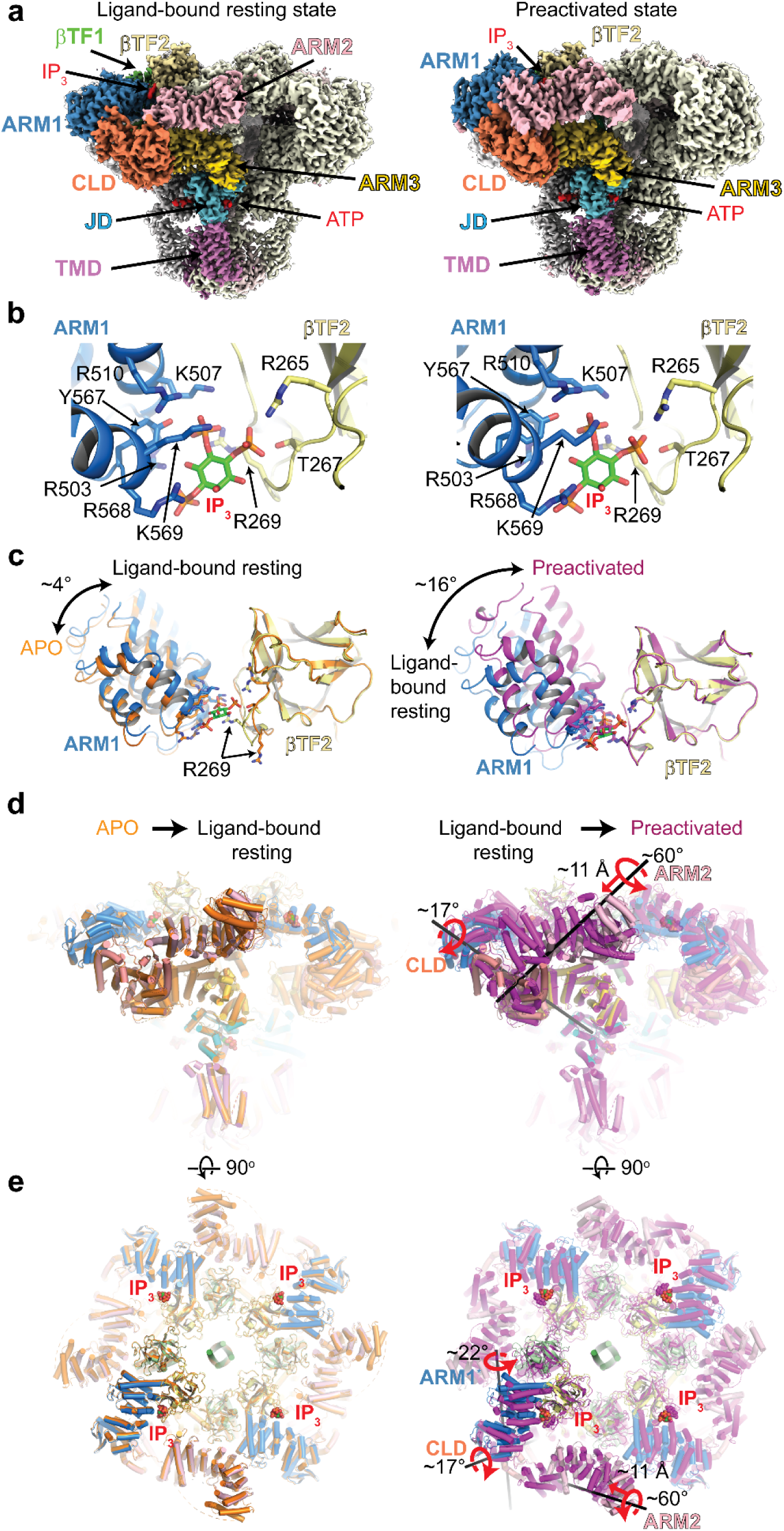
Structures of hIP_3_R-2 in the presence of IP_3_ and ATP. **a,** Composite cryo-EM maps of hIP_3_R-2 in ligand-bound resting (left) and preactivated (right) states. Domains in one subunit are colored as in Fig. 1. Densities for IP_3_ and ATP are shown in red and labeled. **b,** Close-up views of the IP_3_ binding sites of hIP_3_R-2 in resting (left) and preactivated (right) states. Residues in contacting IP_3_ are shown as sticks. **c,** Comparison of the IP_3_ binding sites between apo (orange) and resting state (blue and yellow for ARM1 and βTF2, respectively) on the left and between the resting and preactivated (magenta) states on the right. Structures are aligned on their βTF2, and the rotation of the ARM1 is indicated by a curved arrow. **d-e,** Ribbon representation of the hIP_3_R-2 structures superposed on their TMD. One subunit is shown in full; the others are semi-transparent. Domains coloring for the ligand-bound resting state is the same as in Fig. 1. The apo and preactivated structures are colored in orange and magenta, respectively. The black rods denote the rotation and translation axes. Red curved and straight arrows indicate the direction of the rotations and translations, respectively, for the labeled domains. In the preactivated ensemble, ARM2 retracts toward the CLD while the neighboring ARM1/βTF2 cleft closes.

Relative to the resting state, the IP_3_-binding network in the preactivated state remains largely unchanged, but ARM1 rotates by an additional ∼16° toward βTF2, accompanying coordinated rearrangements across the cytosolic assembly when aligned on the TMD (Fig. 3b-e). The most prominent change involves ARM2. In the apo and resting states, ARM2 adopts an extended conformation in which it forms extensive contacts with ARM1 of the adjacent subunit (Fig. 3d-e). In contrast, the preactivated state ARM2 undergoes a rigid-body rotation of ∼60° and 11 Å translation that brings it closer to the ARM1/CLD of the same subunit, yielding a retracted conformation (Fig. 3d-e). This rearrangement reduces intersubunit contacts between ARM2 and ARM1 and strengthens intrasubunit interactions near the ARM1/CLD interface. These changes likely prime the channel for opening, consistent with the assignment of this state as preactivated.

3D classifications revealed deviations from fourfold symmetry, indicating non-uniform switching across subunits (Supplementary Figs. 4-8). In the resting state, all four subunits adopt an identical conformation: ARM2 is in the extended position, forming intersubunit contacts, and the ARM1/βTF2 cleft surrounding bound IP_3_ is not fully closed. In the preactivated state, all four subunits again converge to a single conformation: ARM2 is retracted, strengthening intrasubunit contacts near ARM1/CLD, and the ARM1/βTF2 cleft is fully closed.

By contrast, the intermediate states are asymmetric, with individual subunits sampling distinct combinations of ARM2 positioning and IP_3_-site closure. For example, in the intermediate state 3 structure, subunit A resembles the resting state (ARM2 extended; ARM1/βTF2 cleft not fully closed) (Fig. 4). Subunit B differs from both resting and preactivated conformations, combining a retracted ARM2 with a cleft that remains partially open. Subunit C matches the preactivated state (ARM2 retracted; cleft fully closed). Subunit D adopts a unique configuration relative to the other three subunits, maintaining an extended ARM2 while the ARM1/βTF2 cleft is fully closed. In the intermediate state 4 structure, the pattern is likewise asymmetric but distinct from intermediate state 3: subunit C adopts a conformation similar to the preactivated state (ARM2 retracted; cleft fully closed), whereas subunit D resembles subunit C from intermediate state 3 (ARM2 retracted; cleft not fully closed). Thus, the intermediate-state structures illustrate that ARM2 retraction and cleft closure can proceed independently within the same subunit. Notably, the position of ARM2 in one subunit correlates with the degree of cleft closure in the adjacent subunit. Retraction of ARM2 in a given subunit is associated with closure of the neighboring ARM1/βTF2 cleft, suggesting intersubunit coupling.

**Figure 4:**
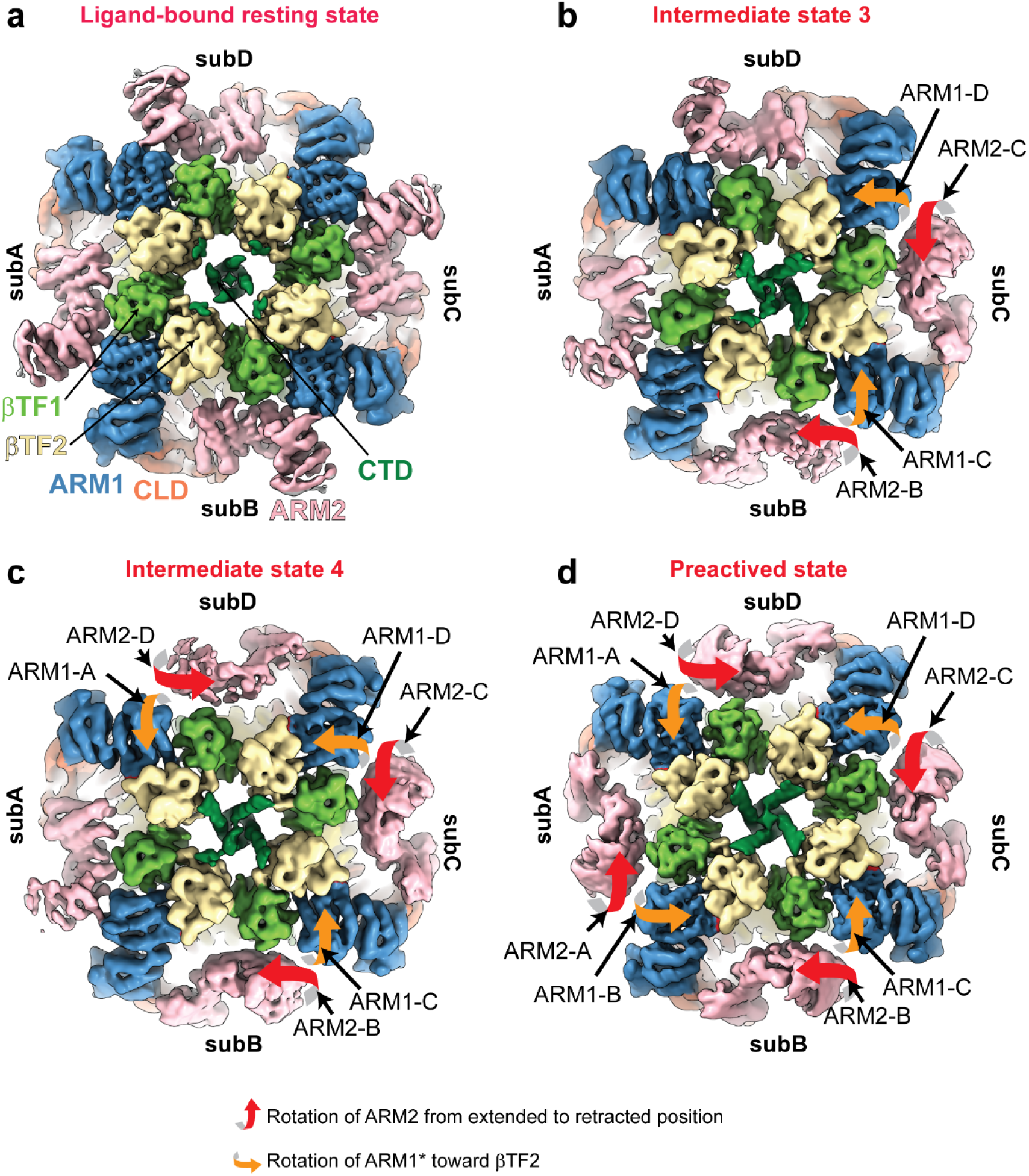
Subunit-asymmetric intermediates in hIP_3_R-2. **a-d,** Consensus cryo-EM maps of hIP_3_R-2 at ligand-bound resting (a), intermediate state 3 (b), intermediate state 4 (c), and preactivated state (d). Maps are low-pass filtered at 5 Å and colored by domains as in Fig. 1. SubA-D denote individual subunits. Orange arrows indicate ARM1 rotation toward βTF2; red arrows indicate ARM2 retraction toward the CLD. Intermediates display subunit-specific combinations of ARM2 positioning and IP_3_-pocket cleft closure, whereas the preactivated ensemble shows a uniform retracted ARM2/closed cleft across all four subunits.

### ATP binding site

All reconstructions from the IP_3_/ATP dataset show strong, well-defined density for ATP bound within the JD, in the ATP-binding pocket previously identified for rIP_3_R-1^29^ and hIP_3_R-3^34,36^(Fig. 5a-b). The adenosine base stacks within a hydrophobic pocket formed by Phe2181, Phe2563, Ile2583, Met2589, and Trp2590, located near the zinc-finger motif, and forms hydrogen bonds with the backbone amide of Phe2563 and the carbonyl group of His2587. The phosphate moieties are coordinated by three basic residues, Arg2174, Lys2127, and Lys2584. Asn2588 is positioned to form hydrogen bonds with the α-phosphate. The ribose ring does not directly interact with protein residues; however, we consistently observe density extending from the ring oxygen across multiple conditions and datasets, plausibly corresponding to a bound water molecule (Fig. 5b).

**Figure 5:**
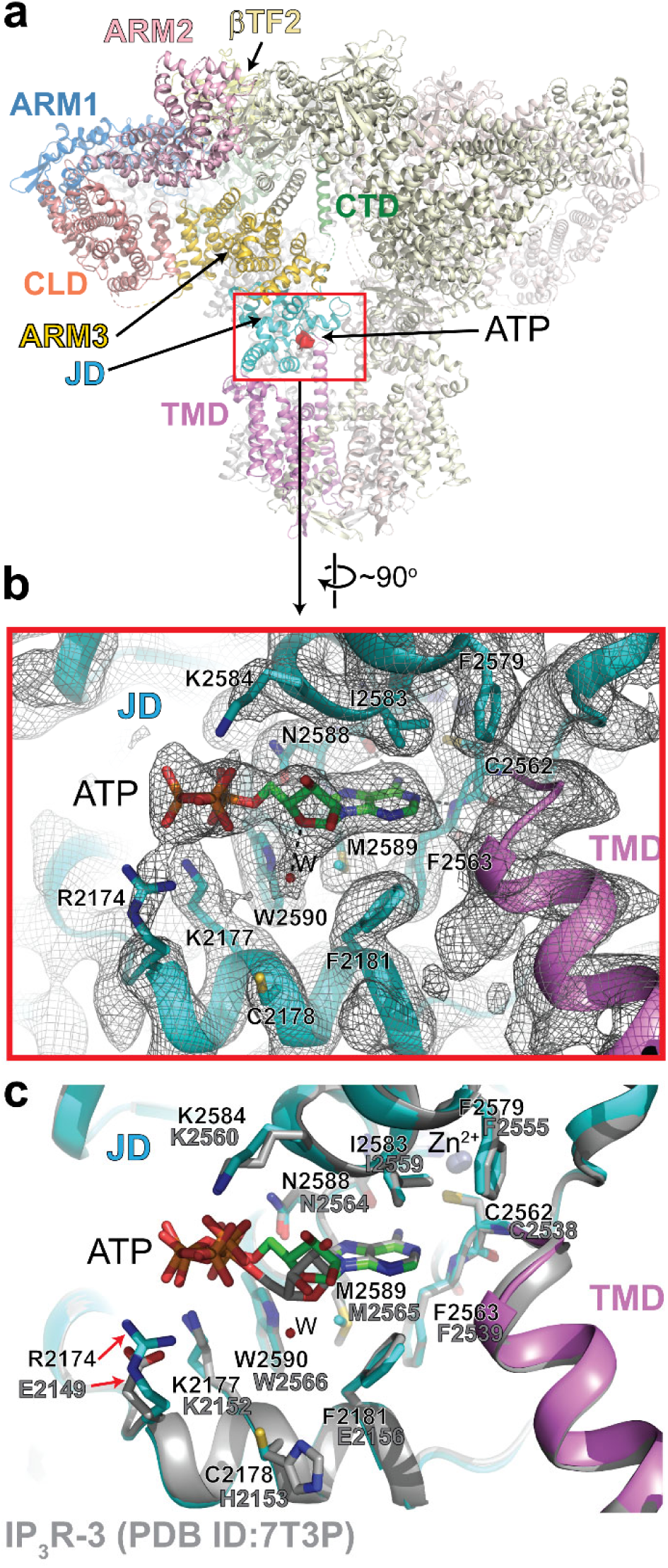
ATP binding to the JD. **a,** Overall structure of hIP_3_R-2 in the ligand-bound resting state, highlighting the ATP binding site in the JD. ATP density is shown in red and labeled. **b,** Close-up view of the ATP binding site with the cryo-EM map (grey mesh). Side chains contacting ATP are shown as sticks, and putative hydrogen bonds are indicated by dashed lines. Arg2174 is well resolved and engages the β-phosphate of ATP; in other states, its density is weaker but remains oriented toward the phosphate moieties. **c,** Comparison of the ATP site in hIP3R-2 (JD, cyan; TMD, magenta) and IP3R-3 (PDB 7T3P, grey), aligned on the JD. Residues are labeled for both subtypes (hIP_3_R-2 above, hIP_3_R-3 below). The substitution of Arg2174 in hIP_3_R-2 for Glu2149 in hIP_3_R-3 may confer additional electrostatic stabilization of the ATP phosphate moieties and contribute to subtype-specific differences in ATP affinity.

IP_3_R-2 has the highest affinity for ATP among IP_3_R subtypes, whereas IP_3_R-3 has the lowest^23–25^. Based on hIP_3_R-3 structures and sequence alignment, we previously proposed that a charge difference in a residue near the phosphate moieties contributes to this disparity^34^. Structural alignment of the JD from hIP_3_R-2 and hIP_3_R-3 shows that the ATP-binding pocket is highly similar and that ATP adopts a comparable pose in both (Fig. 5c). As expected, the most notable difference is Arg2174 in hIP_3_R-2, which corresponds to Glu2149 in hIP_3_R-3. In the ligand-bound resting state, the side chain of Arg2174 is clearly resolved and engages the β-phosphate of ATP. In other structures, the side chain is less well defined but remains oriented toward the phosphate moieties. These observations support our prior proposal that this residue difference partly explains the higher ATP affinity of IP_3_R-2 relative to IP_3_R-3.

### The structure of hIP_3_R-2 in an inactive-like state

In both the apo and IP_3_/ATP datasets, a substantial fraction of particles fall into a loose class in which cytosolic intersubunit contacts are weakened or lost. This architecture parallels the hIP_3_R-3 conformation that increases with Ca^2+^ and predominates at inhibitory concentrations (∼2 mM)^34–36^, and we therefore refer to it as inactive-like. No Ca^2+^ was intentionally added prior to grid preparation in either condition; however, residual free Ca^2+^ in thin films cannot be excluded (see Methods).

In the IP_3_/ATP/Ca^2+^ dataset, the vast majority of particles adopted the loose conformation, with only a small fraction populating compact class, yielding a low-resolution map insufficient for high-resolution refinement (Supplementary Figs. 9-10). The loose class particles segregated into two subclasses: class 1 closely resembles the loose class observed without added Ca^2+^, whereas class 2 exhibits additional features that will be discussed in the next section. We did not observe pore dilation consistent with an open state in either subclass.

To assess the functional state represented by the loose classes, we compared the structures in the loose class from the IP_3_/ATP and IP_3_/ATP/Ca^2+^ (class 1) datasets (Fig. 6). Consensus maps from the global refinement appear practically identical (Fig. 6a). The JDs and TMDs are well defined. In contrast, resolution decreases markedly for the remainder of the protein. Cytosolic domains are resolved in two subunits and largely invisible in the other two, indicating their increased conformational heterogeneity. Local refinement improved the map quality for visible cytoplasmic domains, but not for the invisible ones (Fig. 6b). Close inspection of the local refinement map from the IP_3_/ATP/Ca^2+^ dataset reveals substantial spherical density at the ARM3-JD interface, corresponding to the activatory Ca^2+^ binding site reported for rIP_3_R-1^29,39^ and hIP_3_R-3^34,36^ (Fig. 6c). The site and coordination are consistent with previous structures. Ca^2+^ is coordinated by Glu1930 and Glu1944 in ARM3, and the backbone carbonyl of Thr2605 in JD, with His1932 and Gln1997 in proximity and potentially involved via water-mediated contacts. Coordination of Ca^2+^ at this site is coupled to a ∼10° clamshell-like closure of JD relative to ARM3 (Fig. 6d). In the IP_3_/ATP dataset, density at the same location is considerably weaker and the presence of Ca^2+^ cannot be confidently assessed (Fig. 6c). However, the ARM3-JD arrangement is nearly identical to that observed in the Ca^2+^-bound structure (Fig. 6d). This could mean that the ARM3 and JD can adopt the closed clamshell conformation in the absence of Ca^2+^. Alternatively, Ca^2+^ may be present, but not visualized in the cryo-EM maps due to weak occupancy or local disorder at the site.

**Figure 6:**
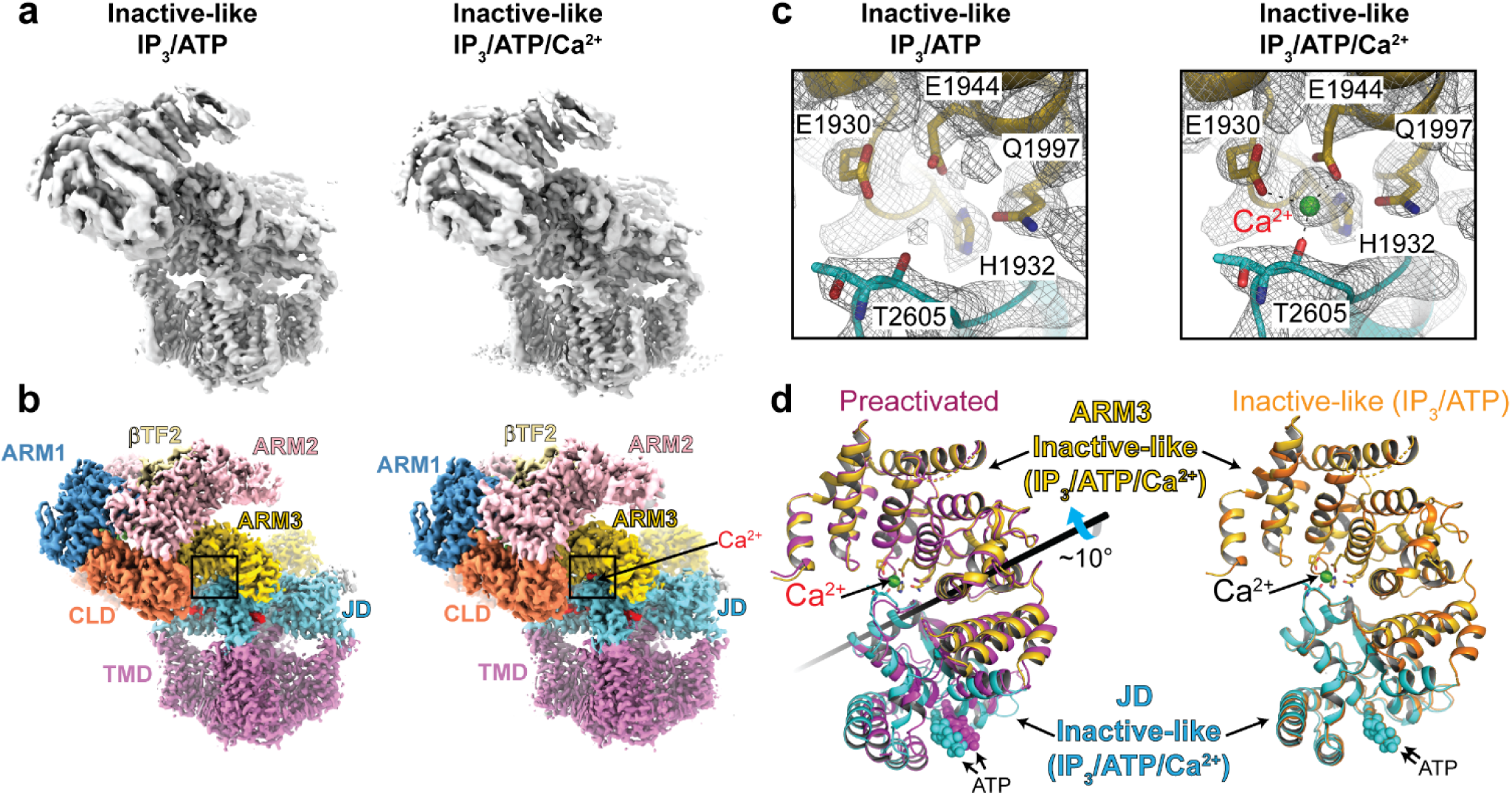
Inactive-like structures of hIP_3_R-2. **a-b,** Unsharpened consensus (**a**) and composite (**b**) maps of the inactive-like structures from IP_3_/ATP (left) and IP_3_/ATP/Ca^2+^ conditions. Composite maps were assembled from locally refined subregions. **c,** Close-up view of the activatory Ca^2+^ binding site at the ARM3-JD interface along with the cryo-EM map (grey mesh) from local refinements from IP_3_/ATP (left) and IP_3_/ATP/Ca^2+^ (right) datasets. In the IP_3_/ATP/Ca^2+^ dataset, a spherical density at the site is consistent with Ca^2+^ occupancy, while no comparable ion density is evident in the IP_3_/ATP dataset, and ion identity is not assigned. **d,** Comparison of ARM3-JD across ensembles: preactivated (magenta), inactive-like IP_3_/ATP (orange), and inactive-like IP_3_/ATP/Ca^2+^ (ARM3, yellow; JD, cyan), aligned on ARM3. The black rod denotes the JD rotation axis relative to ARM3.

A second Ca^2+^ binding site at the CLD-ARM2 interface has been reported in Ca^2+^-inhibited structures of hIP_3_R-3^35,36^, and mutational analysis suggests that this site contributes, at least in part, to Ca^2+^-dependent inhibition of hIP_3_R-3 channels^40^. In our hIP_3_R-2 maps, we do not observe density that can be attributed with high confidence to a bound Ca^2+^ ion at the analogous CLD-ARM2 site or elsewhere.

### hIP_3_R-2 clustering

About half of the particles from the IP_3_/ATP/Ca^2+^ dataset (class 2) exhibited substantial density adjacent to a complete hIP_3_R-2 tetramer (Supplementary Fig. 9). Inspection of this additional density indicated that it corresponds to a second hIP_3_R-2 channel interacting with the first. To define the nature of this interaction, we re-extracted the particles with a larger box size to accommodate both channels (Fig. 7a and Supplementary Fig. 11). Despite the limited resolution, the resulting map clearly revealed two hIP_3_R-2 tetramers interacting through their cytosolic domains with a twofold symmetry (Fig. 7a). Rigid body fitting of the structural models indicates that the channels forming the complex are in loose conformation. In this arrangement, the TMDs of both channels align well on a nearly planar surface, consistent with a geometry compatible with the ER membrane.

**Figure 7:**
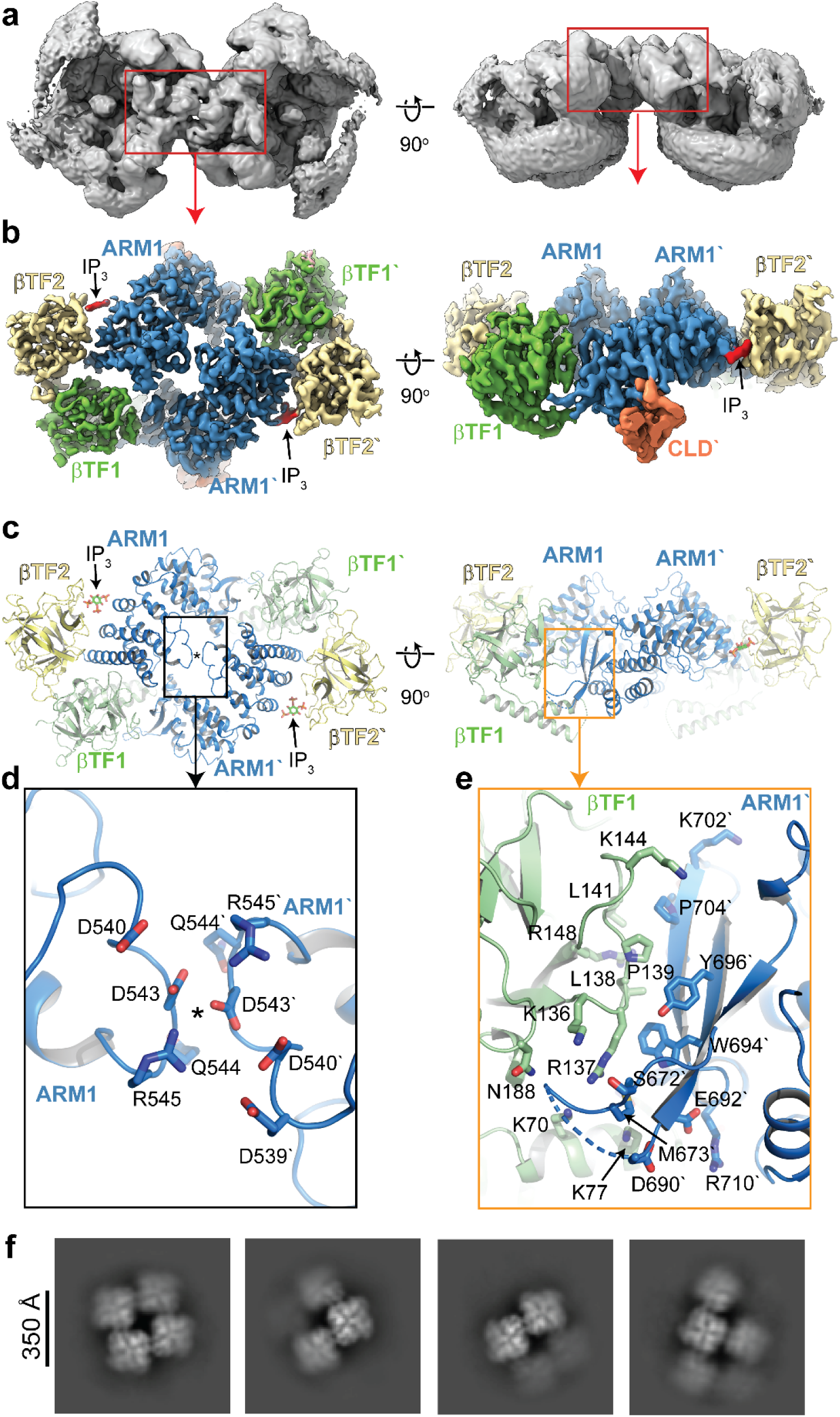
Clustering interface of hIP_3_R-2. **a,** Consensus cryo-EM map of hIP_3_R-2 dimer viewed from the cytosol (left) and along the membrane plane (right). The TMDs are nearly coplanar with a small tilt relative to a common membrane plane, consistent with ER membrane geometry. Red boxes indicate the viewed regions in panel **b-e**. Rigid-body fitting indicates both channels adopt the loose conformation. **b-c,** Locally refined cryo-EM map (**b**) and ribbon representation (**c**) of the domains forming the dimerization interface, viewed as in panel a. Domains are colored as in Fig. 1. Asterisk (*) denotes the approximate two-fold symmetry axis. **d-e**, Close up view of the boxed regions in panel c. Select residues are shown as sticks and labeled. Labels with a prime (′) denote residues from the partner channel. **f,** Representative 2D class averages revealing higher-order assemblies involving three or more channels. These classes support a shared interface but did not yield 3D reconstructions in the current dataset.

Although the dimer map was not sufficiently resolved for de novo model building across the entire assembly, the initial reconstruction generated with the smaller box size provided a substantially better map for one subunit and for the domains of the second channel that form the inter-channel interface (Supplementary Figs. 9 and 11). To enhance local map quality at the putative clustering interface, we carried out focused refinements with masks spanning ARM1, βTF1, and βTF2. This yielded a local resolution of ∼3.5 Å at the interface, permitting side-chain assignment for most residues (Fig. 7b and Supplementary Fig. 11)

The two channels interact through their ARM1 and βTF1 domains with approximate twofold symmetry (Fig. 7b-c). Near the symmetry axis, the loops comprising residues 540-545 from both subunits are in close proximity, but their contribution to the stability of the complex appears limited, as they bury about 10% of the total interfacial area (Fig. 7c-d). The interaction is predominantly hydrophilic, involving Asp540, Asp543, Gln544, and Arg545. The dominant contacts flank this central region and are formed by ARM1 on one channel engaging βTF1 on the other, accounting for the remainder of the interface (Fig. 7c and e). In total, the interface buries 2,818 Å^2^ (∼1,409 Å^2^ per channel), as computed with NACCESS^41^ (probe radius 1.4 Å). The three-stranded β-sheet within ARM1 (residues 667-713) interacts with the loop comprising residues 135-144 on βTF1, forming the core of the interface. Within the β-sheet, the loop comprising residues 676-689 is not resolved in the cryo-EM maps, suggesting higher flexibility, but they may also contribute to the complex formation. Additional contacts within this region are observed through the two α-helices extending from the βTF1 toward the ARM1 (Fig. 7c and e). This region is less well defined, and residues 79-85 are completely invisible in the maps.

While 3D analysis revealed a dimeric arrangement, 2D classifications indicate higher-order clustering of hIP_3_R-2 involving three or more channels (Fig. 7e). It was not possible to obtain 3D maps for these clusters with the current data. However, the 2D classes clearly indicate that these higher-order clusters are mediated through the same mode of interactions observed in the dimeric structure. Importantly, the TMDs of these channels in the assemblies are aligned on a nearly planar surface, indicating that a similar mode of assembly is plausible on the ER membrane.

## DISCUSSION

Here, we determined the cryo-EM structures of hIP_3_R-2 across multiple ligand conditions, defining compact (apo, resting, and preactivated), loose (inactive-like), and asymmetric intermediates. These structures delineate IP_3_ and ATP recognition and support a mechanism in which ARM2 retraction is coupled to closure of the adjacent ARM1/βTF2 cleft. We also observe ligand-dependent dimerization and higher-order clustering mediated by ARM1/βTF1 contacts with nearly planar alignment of TMDs, consistent with the endoplasmic reticulum geometry.

hIP_3_R-2 shares the core architecture of rIP_3_R-1 and hIP_3_R-3, with a modest isoform-specific tilt of ARM2 in the apo state compared to hIP_3_R-3 (Fig. 2). The IP_3_-binding sites in all three subtypes are highly similar and undergo similar rearrangements upon IP_3_ binding^29,30,34–36^. Likewise, global conformational changes upon ligand binding are consistent with a conserved mechanism for activation and gating. At the same time, physiological differences among the subtypes likely arise from subtle distinctions that shift ligand sensitivity and the kinetics of conformational transitions. One such difference is observed at the ATP binding, where the presence of Arg2174 in hIP_3_R-2 versus Glu2149 in hIP_3_R-3 provides additional electrostatic stabilization/coordination of the phosphate moiety of ATP, potentially yielding tighter binding (Fig. 5c). hIP_3_R-2 also has the highest affinity for IP_3_; however, our analysis did not identify a specific structural feature that explains this difference.

Inspired by our previous work on hIP_3_R-3 reporting SBP engagement at the IP_3_-binding site^33^, we performed focused 3D classification of the apo hIP_3_R-2 IP_3_-binding pocket to assess whether a similar interaction occurs. Our analysis reveals that the SBP adopts distinct conformations, involving one that occupies the IP_3_-binding site. Unlike hIP_3_R-3, hIP_3_R-2 also shows that SBP interacts with the adjacent subunits. These observations suggest that the SBP is dynamically involved in intra- and intersubunit interactions, which may contribute to the regulation of channel activity. Identifying regulatory elements that bias SBP conformations will be an important step toward establishing their mechanistic role in IP_3_R activity.

Cryo-EM reconstructions of hIP_3_R-2 in the presence of IP_3_ and ATP reveal conserved IP_3_ recognition across distinct protein conformational states (Fig. 3). In a ligand-bound resting state, IP_3_ induces local changes, including a modest clamshell motion (ARM1 rotates ∼4° toward βTF2), and the overall architecture resembles apo. In the preactivated state, ligand contacts are preserved, but the cytosolic assembly rearranges substantially: ARM2 retracts toward the CLD and the adjacent ARM1/βTF2 cleft fully closes with an additional ∼16° ARM1 rotation. Asymmetric intermediates show that ARM2 retraction and cleft closure can occur independently within subunits, while ARM2 retraction in one subunit correlates with closure of the neighboring cleft, indicating intersubunit coupling. The coexistence of resting, preactivated, and asymmetric intermediates under saturating IP_3_ suggests that additional determinants modulate the conformational equilibrium.

IP_3_-induced rearrangements in hIP_3_R-2 closely parallel those reported for rIP_3_R-1^29,30^ and hIP_3_R-3^34–36^, suggesting a conserved trajectory from resting to preactivated toward pore opening. In hIP_3_R-3, Ca^2+^ binding at the activating ARM3-JD interface correlates with pore opening^34,36^. Although we did not capture a clearly dilated pore in hIP_3_R-2, the similarity of IP_3_-driven rearrangements supports a comparable pathway. In the IP_3_/ATP/Ca^2+^ dataset, we observe density at the activating ARM3-JD site consistent with Ca^2+^ occupancy; nevertheless, these particles adopt a loose, inactive-like conformation. Unlike hIP_3_R-3^35,36^, we did not detect density for additional Ca^2+^-binding sites in these inactive-like structures. Absence of detectable density does not necessarily imply absence of binding, as Ca^2+^ density can be obscured by imperfect coordination or local flexibility. Accordingly, it remains uncertain whether the loose structures represent desensitized or Ca^2+^-inhibited states, and we conservatively refer to them as inactive-like, pending further structural or functional evidence.

IP_3_Rs self-assemble into discrete groups in a ligand-dependent (IP_3_ and Ca^2+^) manner and exhibit distinct physiological properties compared to individual receptors^37,42–45^. These clusters lead to Ca^2+^ puffs, which are localized, transient ER Ca^2+^ release events arising from coordinated channel opening^28,37^. Our structural observations offer clues into the mechanism of such ligand-dependent clustering (Fig. 7). The interchannel interface appears to be accessible upon conformational changes induced by IP_3_ and Ca^2+^. Consistently, the 2D class averages indicate array-like arrangements of two or more channels on a nearly planar surface with aligned transmembrane domains, suggesting that this mode of interaction permits clustering without major geometric constraints and is plausible on the ER membrane. This higher-order arrangement would be most compatible with ordered cytosolic assemblies in all four subunits, which we do not observe in the inactive-like structures. Another consideration is that clustering has been reported to dampen Ca^2+^-dependent inhibition and promote concerted behavior, increasing Ca^2+^ release^43,44^. This may indicate that the structures captured represent an intermediate state en route to fully coordinated clusters, with additional conformational changes and/or higher Ca^2+^ required.

## METHODS

### Construct design and cloning

The full-length hIP_3_R-2 coding sequence (ITPR2; UniProt Q14571) was codon-optimized for expression in *Spodoptera frugiperda* (Sf9) cells and chemically synthesized (Gene Universal Inc.) as five overlapping double-stranded DNA fragments (1,966, 1,958, 1,957, 1,958, and 425 bp). Fragments were assembled by overlap-extension PCR using primers annealing to the 5′ end of fragment 1 and the 3′ end of fragment 5 to yield the complete open reading frame (ORF). The assembled ORF was fused at its C terminus to a thrombin cleavage site and a Twin-Strep tag, and subcloned into the pACEBac1 vector using BamHI and XbaI. A Kozak consensus sequence was incorporated immediately upstream of the start codon to enhance expression in Sf9 cells. The accuracy of the final constructs was verified by Sanger sequencing before being incorporated into baculovirus using the MultiBac expression system and baculovirus preparation.

### Protein expression and purification

Sf9 cells at 3×10^6^ cells/mL were infected and harvested 48 h post-infection by centrifugation (2,000×g, 10 min, 4 °C). Pellets were washed once in PBS, collected again (3,000×g, 10 min, 4 °C), flash-frozen in liquid nitrogen, and stored at −80 °C.

For purification, frozen pellets were thawed on ice and resuspended in lysis buffer (Buffer A: 200 mM NaCl, 40 mM Tris-HCl pH 8.0, 5 mM EDTA pH 8.0, 2 mM DTT) supplemented with 1 mM PMSF. Cells were disrupted using an Avestin EmulsiFlex-C3. The lysate was clarified at 7,000×g for 10 min, and membranes were collected by ultracentrifugation at 185,000×g for 1 h (Beckman Coulter Type 45 Ti rotor). Membrane pellets were homogenized in ice-cold buffer A and solubilized with 0.5% lauryl maltose neopentyl glycol (LMNG; Anatrace, cat. NG310) and 0.1% glycodiosgenin (GDN; Anatrace, cat. GDN101) at ∼100 mg/mL membrane concentration for 4 h with gentle mixing. Insoluble material was removed by ultracentrifugation as above, and the supernatant was applied by gravity to Strep-Tactin XT 4Flow resin (IBA Lifesciences). The resin was washed with 20 column volumes of buffer A containing 0.5% CHAPS (Anatrace, cat. C316S), and bound protein was eluted in the same buffer supplemented with 100 mM D-biotin (pH 8.0). Peak fractions containing hIP_3_R-2 were pooled and polished by size-exclusion chromatography (SEC) on a Superose 6 Increase 10/300 GL column (Cytiva) equilibrated in SEC buffer (200 mM NaCl, 20 mM Tris-HCl pH 8.0, 5 mM EDTA pH 8.0, 2 mM DTT, 0.5% CHAPS). The sample was used immediately for nanodisc preparation.

The membrane scaffold protein MSP1E3D1 expression vector (pMSP1E3D1; Addgene plasmid # 20066; http://n2t.net/addgene:20066; RRID: Addgene_20066), was a gift from Stephen Sligar^46^. MSP1E3D1 was produced in *E. coli* BL21(DE3) cells and purified by Ni^2+^-chelate affinity chromatography as described previously^47^. The histidine tag was removed by TEV protease digestion, and the protein was further purified by SEC using a HiLoad 16/600 Superdex 200 pg column (Cytiva) equilibrated with 300 mM NaCl and 40 mM Tris-HCl, pH 8.0. Peak fractions were pooled, aliquoted, flash-frozen in liquid nitrogen, and stored at -80°C.

### Nanodisc incorporation of hIP_3_R-2

1,2-dioleoyl-sn-glycero-3-phosphocholine (DOPC), dissolved in 1% CHAPS, was mixed with hIP_3_R-2 detergent micelles at a lipid-to-protein molar ratio of 200:1 and incubated for 30 min at 4 °C with gentle agitation. MSP1E3D1 was then added at a molar ratio of 4:1 (MSP1E3D1:hIP_3_R-2), and the mixture was incubated for an additional hour at 4 °C. The detergent was removed by dialysis overnight at 4 °C against 2 L of SEC buffer lacking detergent (200 mM NaCl, 20 mM Tris-HCl pH 8.0, 5 mM EDTA pH 8.0, 2 mM DTT), allowing nanodisc self-assembly. hIP_3_R-2 nanodiscs were isolated by SEC on a Superose 6 Increase 10/300 GL column (Cytiva) equilibrated in the same detergent-free buffer. Fractions corresponding to hIP_3_R-2 nanodiscs were pooled and concentrated to ∼4 mg/mL using a 100 kDa centrifugal filter (Millipore), clarified by ultracentrifugation (260,000×g, 10 min; ThermoFisher S110AT rotor). The concentration typically dropped to ∼2.5 mg/mL.

### Planar lipid bilayer electrophysiology

Single-channel recordings of hIP3R-2 were performed using a MECA-4 chips with 100 µm cavities (Ionera Technologies) on an Orbit Mini system equipped with a temperature control unit (Nanion Technologies)^48^. Planar lipid bilayers were formed using the air-bubble technique from a DOPC/DOPE mixture (5:3 molar ratio, Avanti Polar Lipids) dissolved in n-decane (Sigma-Aldrich) to a final concentration of 20 mg/mL. Chips were filled with 150 µL of symmetrical recording buffer containing 20 mM HEPES (pH 7.4), 150 mM KCl, 2 mM TCEP, 1 mM ATP, 0.5 mM MgCl_2_, 10 µM IP_3_, 5 mM EGTA, and varying CaCl_2_ to reach desired free Ca^2+^ concentrations. Cavities were wetted prior to bilayer formation. A lipid-coated pipette tip was used to form an air bubble beneath the buffer surface near a cavity. The bubble was slowly expanded and retracted to paint the lipid monolayer. Bilayers were ruptured and reformed several times until a single, stable bilayer was achieved, which was confirmed by monitoring its capacitance and conductance. Channel incorporation was performed using hIP_3_R-2 nanodics following an approach described in ^49,50^. 0.5 µL of hIP_3_R-2 reconstituted in nanodiscs (20 µg/mL in storage buffer containing 20 mM HEPES pH 7.4, 150 mM KCl, 2 mM TCEP, 5 mM EGTA, and 2.5% glycerol) was added near the cavities. The solution was gently mixed, and a holding potential of +40 mV was applied across the membrane to promote fusion. Free Ca^2+^ concentrations were calculated using MaxChelator and verified using the fluorescent indicator, Fluo-8 sodium salt (AAT Bioquest). Experiments were repeated using protein from at least three different purifications for each condition tested.

All recordings were performed at 22 °C under a holding potential of +60 mV. Signals were acquired at a sampling rate of 10 kHz and a final bandwidth of 1 kHz. Single-channel analysis, including the calculation of open probability (Po), was conducted using Clampfit (v10.7). For analysis and presentation, currents were digitally filtered at 500 Hz and 200 Hz, respectively^51^.

### Cryo-EM sample preparation and data collection

Apo hIP_3_R-2 nanodiscs (∼2.5 mg/mL) were supplemented immediately prior to vitrification with 0.025% (w/v) fluorinated Fos-Choline-8 (FC-8; Anatrace, Anagrade; cat. F300F). For IP_3_/ATP and IP3/ATP/Ca^2+^ samples, hIP3R-2 nanodiscs were supplemented with 50 μM IP_3_ (from a 10 mM stock in water), 1 mM ATP (from a 100 mM stock, pH 7.2), either without CaCl_2_ (IP_3_/ATP) or with 4 mM CaCl_2_ (IP_3_/ATP/Ca^2+^), followed by the addition of 0.025% (w/v) FC-8 immediately prior to plunging. In the presence of 5 mM EDTA, the calculated free Ca^2+^ for the IP_3_/ATP/Ca^2+^ mixture is ∼10 nM (MaxChelator). We note that the free Ca^2+^ during grid preparation may deviate from calculated values due to trace Ca^2+^ leaching and the limited buffering capacity in the small volumes applied to grids.

For grid preparation, 2.5 μL of sample was applied to Quantifoil R1.2/1.3 Cu 300 mesh grids (Electron Microscopy Sciences) glow-discharged for 30 s at 25 mA. Grids were blotted for 4 s at blot force 10 using two layers of PELCO 595 filter paper (Ted Pella, prod. no. 47000-100) and plunge-frozen into liquid ethane with a Vitrobot Mark IV (Thermo Fisher) set to 8 °C and 100% humidity. Filter papers were not pretreated with Ca^2+^ chelators or other chemicals. Optimization of the plunging conditions was performed by imaging the grids using Glacios microscope (Thermo Fisher).

All data collection was performed using Titan Krios G4i (Thermo Fisher) equipped with a BioQuantum K3 detector at Vanderbilt cryo-EM facility. Movies were recorded as 50-frame exposures at a nominal magnification of 105,000×, yielding a calibrated pixel size of 0.822 Å/pixel using the automated imaging software EPU (Thermo Fisher). The total dose was 54.4-55.5 e^−^/Å^2^. Four to five shots per hole were acquired, and defocus values ranged from −0.8 to −2.2 μm. Apo and IP_3_/ATP/Ca^2+^ datasets were collected from single grids, whereas the IP_3_/ATP dataset was collected from three grids.

### Cryo-EM data processing

All image processing was performed using CryoSparc (v4.6.2 to v4.7.1)^52^. Motion correction and CTF estimations were performed locally using Patch Motion Correction and Patch CTF Estimation. Micrographs with poor image properties (thin or thick ice, CTF fit resolution worse than 6 Å, large drift) were excluded. Initial particle picking was performed by blob search and particles were then binned 4x and extracted (Supplementary Figs. 1, 4, and 9). After 2D classification, classes with clear structural features were re-extracted at full size (512px). Ab initio reconstructions with 4 classes (apo) or 6 classes (IP_3_/ATP and IP_3_/ATP/Ca^2+^) were performed, followed by heterogeneous refinement. High-quality particles from these reconstructions were used to train a Topaz picker^53^ (200 micrographs), and a new particle set was selected using Topaz^53^ (v0.2.5a) and processed as above. This process was repeated three times to maximize particle yield. All particles from each iteration were merged into a single set after removing the duplicate particles, re-extracted at full size (512px), and subjected to a final heterogeneous refinement. Classes with unique features are grouped and refined using non-uniform (NU) refinement^54^. After refinement, reference-based motion correction^55^ and Local CTF refinement^56^ were performed, followed by another round of NU refinement (Supplementary Figs. 1, 4, and 9).

Under apo conditions, particles segregated into two major classes: compact and loose (Supplementary Fig. 1). The compact class, refined with C4 symmetry, yielded a reconstruction at 2.95 Å average resolution that we designate as the apo state. The map quality for the loose class was considerably lower, precluding model building. Nevertheless, its overall architecture is consistent with an inactive-like conformation.

To assess the SBP occupancy and conformations, we applied C4 symmetry expansion to the apo particle stack, subtracted density for all regions except the N-terminal cytosolic assembly of a single subunit (βTF1, βTF2, ARM1, ARM2, and CLD), and carried out focused 3D classification without angular or translational alignment using a soft mask encompassing the IP_3_-binding site. Class volumes were low-pass filtered to 5 Å for visualization (Supplementary Fig. 3).

In the IP_3_/ATP dataset, particles in the compact class exhibit structural heterogeneity (Supplementary Fig. 4). To separate the particles, we first performed 3D classification without angular or translational alignment using a mask encompassing the entire cytosolic domains (Supplementary Fig. 4). The resulting classes fell into three major groups: resting (with all four ARM2s extended), preactivated (all four ARM2s retracted), and intermediate (mixed ARM2 configurations). To further resolve the intermediate particles, we prepared four masks encompassing the ARM2s of each subunit and performed 3D classification for each mask separately. We then regrouped the particles into all possible ARM2 arrangements. Some classes lacked interpretable ARM2 density, suggesting higher flexibility, and were excluded from further analysis. Particles with four extended or four retracted ARM2s were merged with the resting or preactivated state classes, respectively. NU-refinement of the intermediate classes revealed 3D reconstructions with varying resolutions. Intermediate states 1 and 2 were worse than 6 Å and were not used for further analysis, although their quality was sufficient to confirm these conformations (Supplementary Fig. 5). Intermediate states 3 and 4 produced better maps that allowed assessment of the relative orientation of individual domains but did not permit unambiguous model building (Supplementary Fig. 5). We also observed heterogeneity in the TMD of the preactivated class. To improve the map quality, we performed focused 3D classification using a mask encompassing TMD and retained particle subset that yielded reconstructions with well-defined TMD density (Supplementary Figs. 4-5). Particles in the loose class from the IP_3_/ATP dataset were subjected to another round of 3D classification to remove the particles with low quality. NU-refinement of the remaining particles, without enforcing any symmetry, resulted in a reconstruction with an average resolution of 3.14 Å (Supplementary Figs. 4).

In the IP_3_/ATP/Ca^2+^ dataset, two major classes, both in loose conformation, were observed (Supplementary Fig. 9). The class 1 particles, refined without symmetry enforcement (C1), yielded a map that is highly similar to the loose-class maps from the other datasets. The second class was also refined in C1. While the main portion of the map resembled class 1, substantial amount of extra density was observed. To determine the nature of this extra density, particles were re-extracted with a larger box size (960 px) and two-fold binned to 480 px (Supplementary Fig. 11). We performed ab initio reconstruction searching for 3 classes and without enforcing symmetry. Following heterogeneous refinement, particles that produced a hIP3R-2 dimer map were then refined with C2 symmetry, resulting in a reconstruction at 4.40 Å resolution.

To improve map quality, we performed local refinements using masks spanning subregions of the consensus reconstructions (Supplementary Figs. 2, 6-8, 10-11). For reconstructions processed with C4 symmetry (apo, resting, and preactivated), we prepared four masks that cover distinct cytosolic domains of one subunit and a mask for the TMD tetramer (Supplementary Figs. 2, 6-7). After symmetry expansion (C4), we performed local refinement for the cytosolic domains. The local refinements for the TMDs were performed without symmetry expansion while enforcing C4 symmetry. For the reconstructions without enforced symmetry (inactive-like (IP_3_/ATP), inactive-like (IP_3_/ATP/Ca^2+^), and dimer), we prepared masks that cover different domains of the entire assembly and performed local refinement using C1 symmetry (Supplementary Figs. 8, 10-11). Local refinement maps were aligned onto the consensus maps in ChimeraX^57^ and merged using the “VOP maximum” command to generate composite maps.

### Model building

Models were built using Coot^58^. We first placed the hIP_3_R-2 model generated using the Alphafold server^59^ into the composite apo map, followed by rigid-body placement of individual domains for a single protomer. Manual rebuilding was then performed, and the protomer was symmetry-expanded (C4) to generate the tetramer. Real-space refinement^60^ using Phenix^61^ was performed with iterative build-refine cycles till a satisfactory model was obtained. Manual building was performed using local refinement maps, while real-space refinements used composite maps. The resulting apo model served as the starting reference for the other datasets, which were processed with the same workflow. Regions lacking interpretable density were omitted. Residues with poorly defined side-chain density were retained with side chains truncated to alanine while preserving residue identity. Coiled-coil segments at the CTD were modeled as poly-alanine based on features visible in unsharpened maps. Validation of the structural models was performed using MolProbity^62^ implemented in Phenix^61^.

### Figure preparation

Figures were prepared using ChimeraX^57^ and The PyMOL Molecular Graphics System (Version 2.0, Schrödinger, LLC).

**Table 1:**
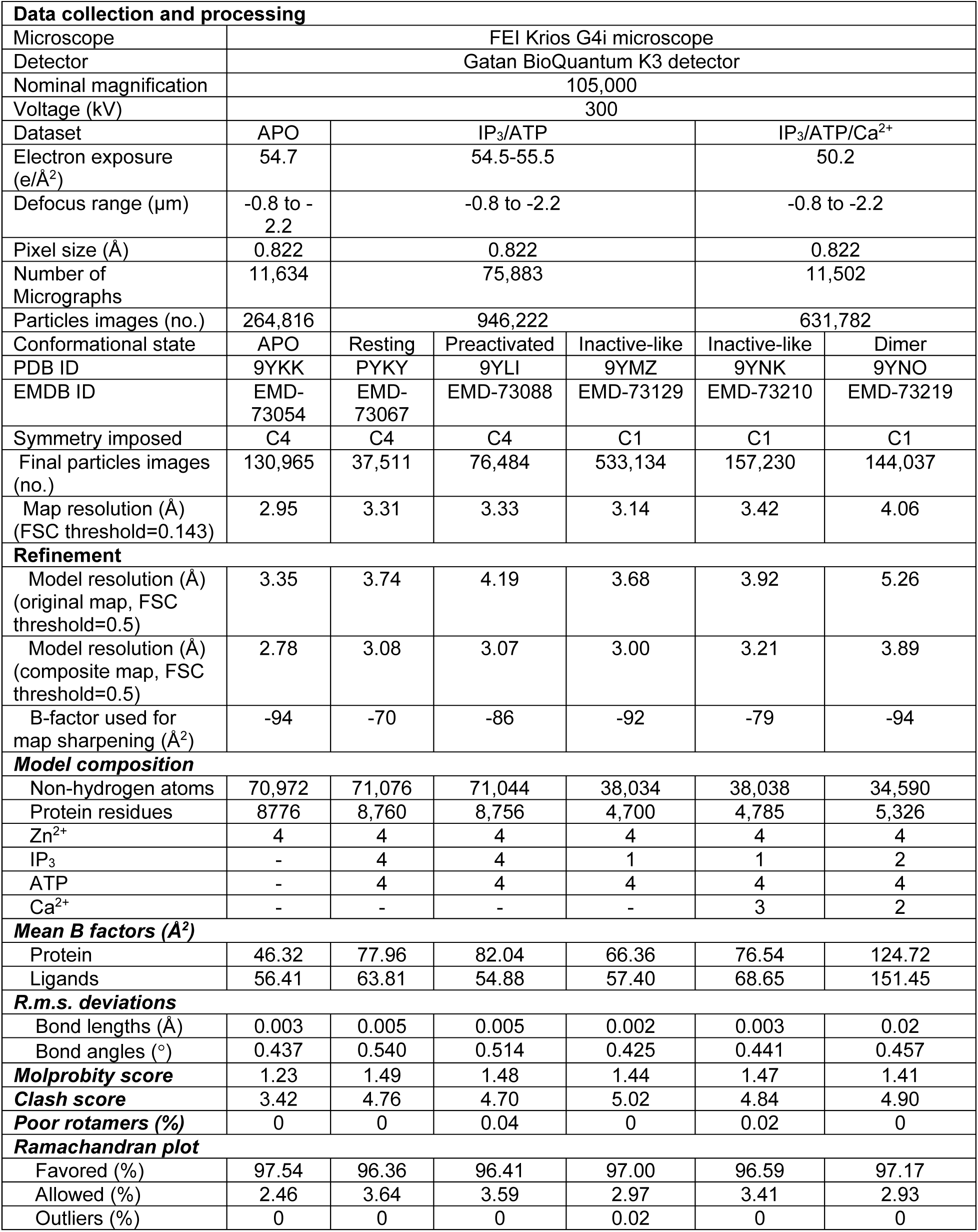
Cryo-EM data collection, refinement, and validation statistics.

|  |  |  |  |  |  |  |
| --- | --- | --- | --- | --- | --- | --- |
| **Data collection and processing** |  |  |  |  |  |  |
| Microscope | FEI Krios G4i microscope |  |  |  |  |  |
| Detector | Gatan BioQuantum K3 detector |  |  |  |  |  |
| Nominal magnification | 105,000 |  |  |  |  |  |
| Voltage (kV) | 300 |  |  |  |  |  |
| Dataset | APO | IP<sub>3</sub>/ATP |  |  | IP<sub>3</sub>/ATP/Ca<sup>2+</sup> |  |
| Electron exposure (e/Å<sup>2</sup>) | 54.7 | 54.5-55.5 |  |  | 50.2 |  |
| Defocus range (μm) | -0.8 to -2.2 | -0.8 to -2.2 |  |  | -0.8 to -2.2 |  |
| Pixel size (Å) | 0.822 | 0.822 |  |  | 0.822 |  |
| Number of Micrographs | 11,634 | 75,883 |  |  | 11,502 |  |
| Particles images (no.) | 264,816 | 946,222 |  |  | 631,782 |  |
| Conformational state | APO | Resting | Preactivated | Inactive-like | Inactive-like | Dimer |
| PDB ID | 9YKK | PYKY | 9YLI | 9YMZ | 9YNK | 9YNO |
| EMDB ID | EMD-73054 | EMD-73067 | EMD-73088 | EMD-73129 | EMD-73210 | EMD-73219 |
| Symmetry imposed | C4 | C4 | C4 | C1 | C1 | C1 |
| Final particles images (no.) | 130,965 | 37,511 | 76,484 | 533,134 | 157,230 | 144,037 |
| Map resolution (Å) (FSC threshold=0.143) | 2.95 | 3.31 | 3.33 | 3.14 | 3.42 | 4.06 |
| **Refinement** |  |  |  |  |  |  |
| Model resolution (Å) (original map, FSC threshold=0.5) | 3.35 | 3.74 | 4.19 | 3.68 | 3.92 | 5.26 |
| Model resolution (Å) (composite map, FSC threshold=0.5) | 2.78 | 3.08 | 3.07 | 3.00 | 3.21 | 3.89 |
| B-factor used for map sharpening (Å<sup>2</sup>) | -94 | -70 | -86 | -92 | -79 | -94 |
| **Model composition** |  |  |  |  |  |  |
| Non-hydrogen atoms | 70,972 | 71,076 | 71,044 | 38,034 | 38,038 | 34,590 |
| Protein residues | 8776 | 8,760 | 8,756 | 4,700 | 4,785 | 5,326 |
| Zn<sup>2+</sup> | 4 | 4 | 4 | 4 | 4 | 4 |
| IP<sub>3</sub> | - | 4 | 4 | 1 | 1 | 2 |
| ATP | - | 4 | 4 | 4 | 4 | 4 |
| Ca<sup>2+</sup> | - | - | - | - | 3 | 2 |
| **Mean B factors (Å<sup>2</sup>)** |  |  |  |  |  |  |
| Protein | 46.32 | 77.96 | 82.04 | 66.36 | 76.54 | 124.72 |
| Ligands | 56.41 | 63.81 | 54.88 | 57.40 | 68.65 | 151.45 |
| **R.m.s. deviations** |  |  |  |  |  |  |
| Bond lengths (Å) | 0.003 | 0.005 | 0.005 | 0.002 | 0.003 | 0.02 |
| Bond angles (°) | 0.437 | 0.540 | 0.514 | 0.425 | 0.441 | 0.457 |
| **Molprobity score** | 1.23 | 1.49 | 1.48 | 1.44 | 1.47 | 1.41 |
| **Clash score** | 3.42 | 4.76 | 4.70 | 5.02 | 4.84 | 4.90 |
| **Poor rotamers (%)** | 0 | 0 | 0.04 | 0 | 0.02 | 0 |
| **Ramachandran plot** |  |  |  |  |  |  |
| Favored (%) | 97.54 | 96.36 | 96.41 | 97.00 | 96.59 | 97.17 |
| Allowed (%) | 2.46 | 3.64 | 3.59 | 2.97 | 3.41 | 2.93 |
| Outliers (%) | 0 | 0 | 0 | 0.02 | 0 | 0 |

## DATA AVAILABILITY

The cryo-EM density maps and atomic coordinates have been deposited in the Electron Microscopy Data Bank (EMDB) and the Protein Data Bank (PDB), respectively, under the following accession numbers.

Apo hIP_3_R-2: PDB 9YKK; EMDB EMD-73048 (consensus), EMD-73054 (composite), and EMD-73049, EMD-73050, EMD-73051, EMD-73052, EMD-73053 (locally refined subregions).

Inactive-like hIP_3_R-2 (ligand-free): EMDB EMD-73225 (consensus).

hIP_3_R-2 in the ligand-bound resting state: PDB 9YKY; EMDB EMD-73061 (consensus), EMD-73067 (composite), and EMD-73062, EMD-73063, EMD-73064, EMD-73065, EMD-73066 (locally refined subregions).

hIP_3_R-2 in the preactivated state: PDB 9YLI; EMDB EMD-73082 (consensus), EMD-73088 (composite), and EMD-73083, EMD-73084, EMD-73085, EMD-73086, EMD-73087 (locally refined subregions).

hIP_3_R-2 in the intermediate state 3: EMDB EMD-73223 (consensus)

hIP_3_R-2 in the intermediate state 3: EMDB EMD-73224 (consensus)

hIP_3_R-2 in the preactivated state: PDB 9YLI; EMDB EMD-73082 (consensus), EMD-73088 (composite), and EMD-73083, EMD-73084, EMD-73085, EMD-73086, EMD-73087 (locally refined subregions).

Inactive-like hIP_3_R-2 (IP_3_/ATP): PDB 9YMZ; EMDB EMD-73111 (consensus), EMD-73129 (composite), and EMD-73112, EMD-73113, EMD-73114, EMD-73115, EMD-73116, EMD-73117, EMD-73118, EMD-73119, EMD-73120, EMD-73121 (locally refined subregions).

Inactive-like hIP_3_R-2 (IP_3_/ATP/Ca^2+^): PDB 9YNK; EMDB EMD-73198 (consensus), EMD-73210 (composite), and EMD-73199, EMD-73201, EMD-73202, EMD-73203, EMD-73204, EMD-73205, EMD-73206, EMD-73207, EMD-73208, EMD-73209 (locally refined subregions).

hIP_3_R-2 dimer (IP_3_/ATP/Ca^2+^): PDB 9YNO; EMDB EMD-73212 (consensus), EMD-73219 (composite), and EMD-73213, EMD-73214, EMD-73215, EMD-73216, EMD-73217, EMD-73218 (locally refined subregions).

hIP_3_R-2 dimer (complete; IP_3_/ATP/Ca^2+^): EMDB EMD-73222 (consensus)

## ACKNOWLEDGEMENTS

EM data collections were conducted at the Center for Structural Biology Cryo-EM Facility at Vanderbilt University. We thank Dr. Melissa Chambers, Dr. Scott Collier, and Mariam Haider for their help in data collection. We acknowledge the use of the Glacios cryo-TEM, which was acquired by NIH award S10 OD030292. We used the DORS storage system supported by the NIH award S10 RR031634. This work was supported by the NIH (R01 GM141251 to E.K.) and Vanderbilt University School of Medicine Basic Sciences.

## AUTHOR CONTRIBUTIONS

E.K. conceived the project, performed cryo-EM data analysis, and wrote the manuscript with input from all authors; C.L. optimized and performed protein expression and purification, performed grid preparation and screening, and model building and refinement; Y.L. performed protein purification and electrophysiology; Q.T. helped with protein purification, grid preparation, screening, and model building and refinement.

## Supplementary Figures

**Supplementary Fig. 1:**
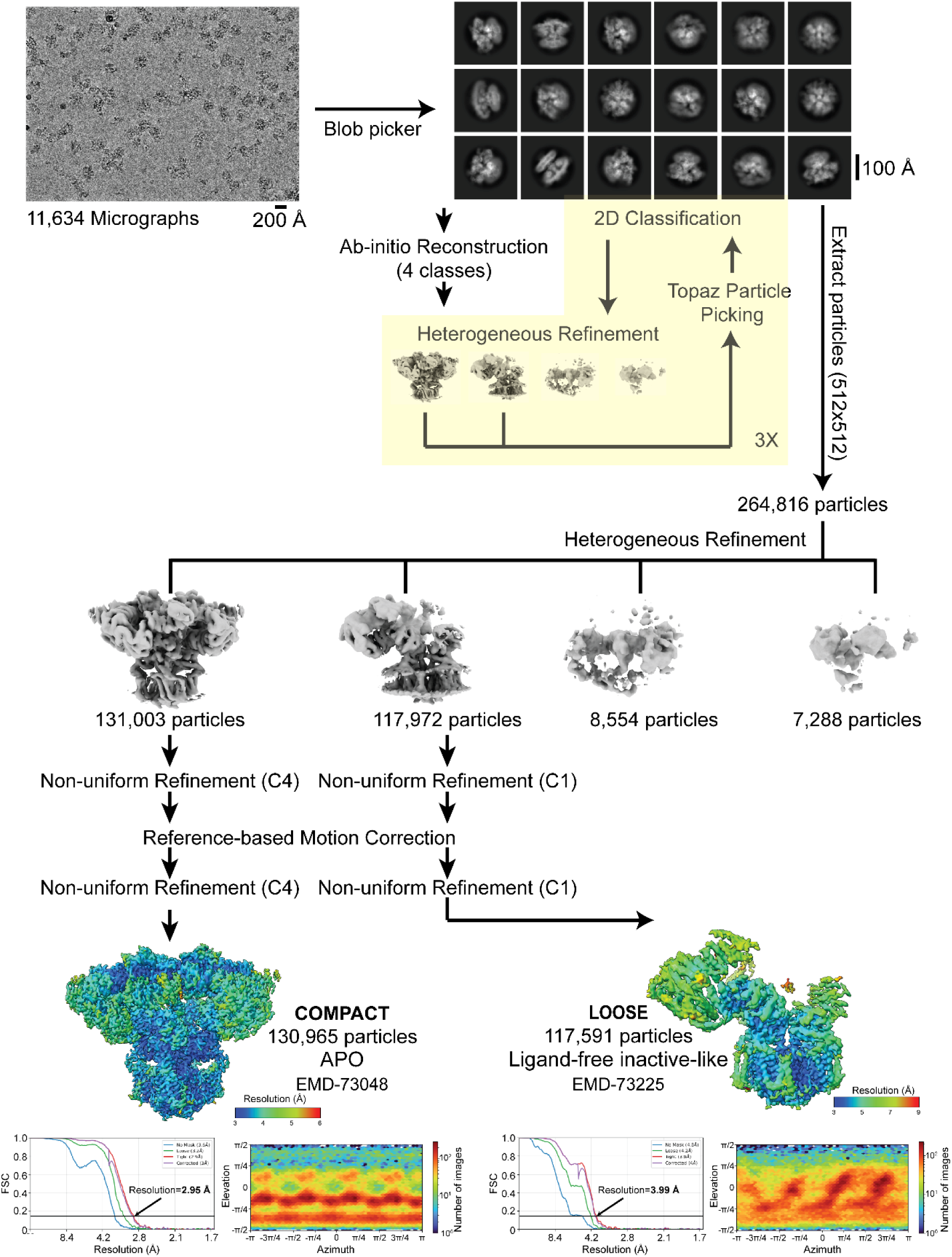
Cryo-EM analysis of hIP_3_R-2 under ligand-free conditions. Representative micrograph of the hIP_3_R-2 dataset, reference-free 2D class averages, and the data-processing workflow from particle picking through 3D classification/refinement, yielding the main classes. Validation plots include local resolution maps, gold-standard FSC curves (0.143 criterion), and particle angular distributions for the principal reconstructions. See the Methods for details on data acquisition and processing.

**Supplementary Fig. 2:**
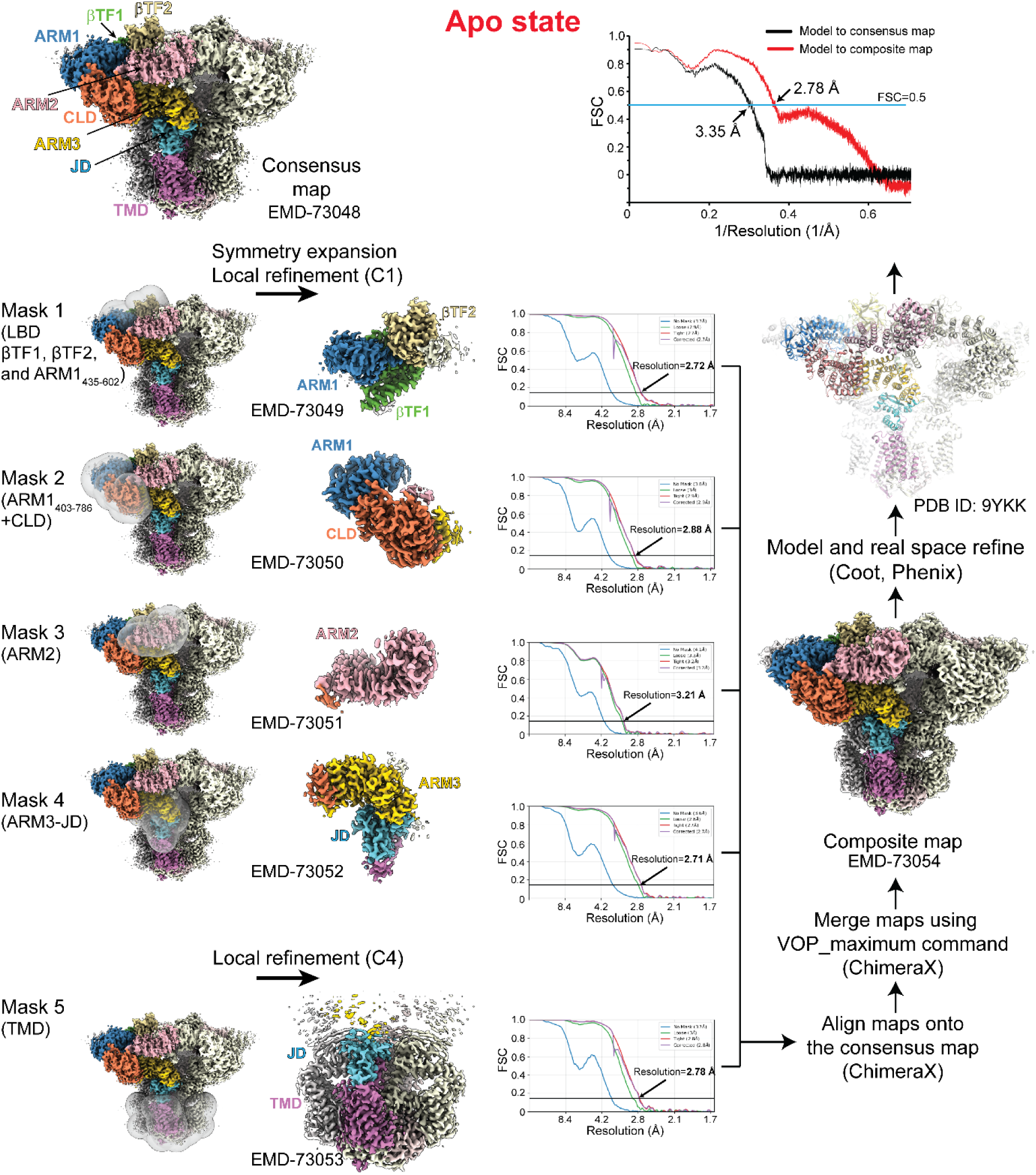
Local refinement of the hIP_3_R-2 apo state. Workflow from the consensus C4-symmetry reconstruction (2.95 Å overall) to symmetry expansion and focused refinements with per-subunit/per-domain masks, followed by assembly of a composite map from local refinements. Masks are shown as transparent overlays; domains are colored as in Fig. 1. Each locally refined map is labeled with its EMDB accession. For each mask, gold-standard FSC curves computed in cryoSPARC (0.143 criterion) are shown. Model-map FSCs compare the refined model against the consensus map (black) and the composite map (red).

**Supplementary Fig. 3:**
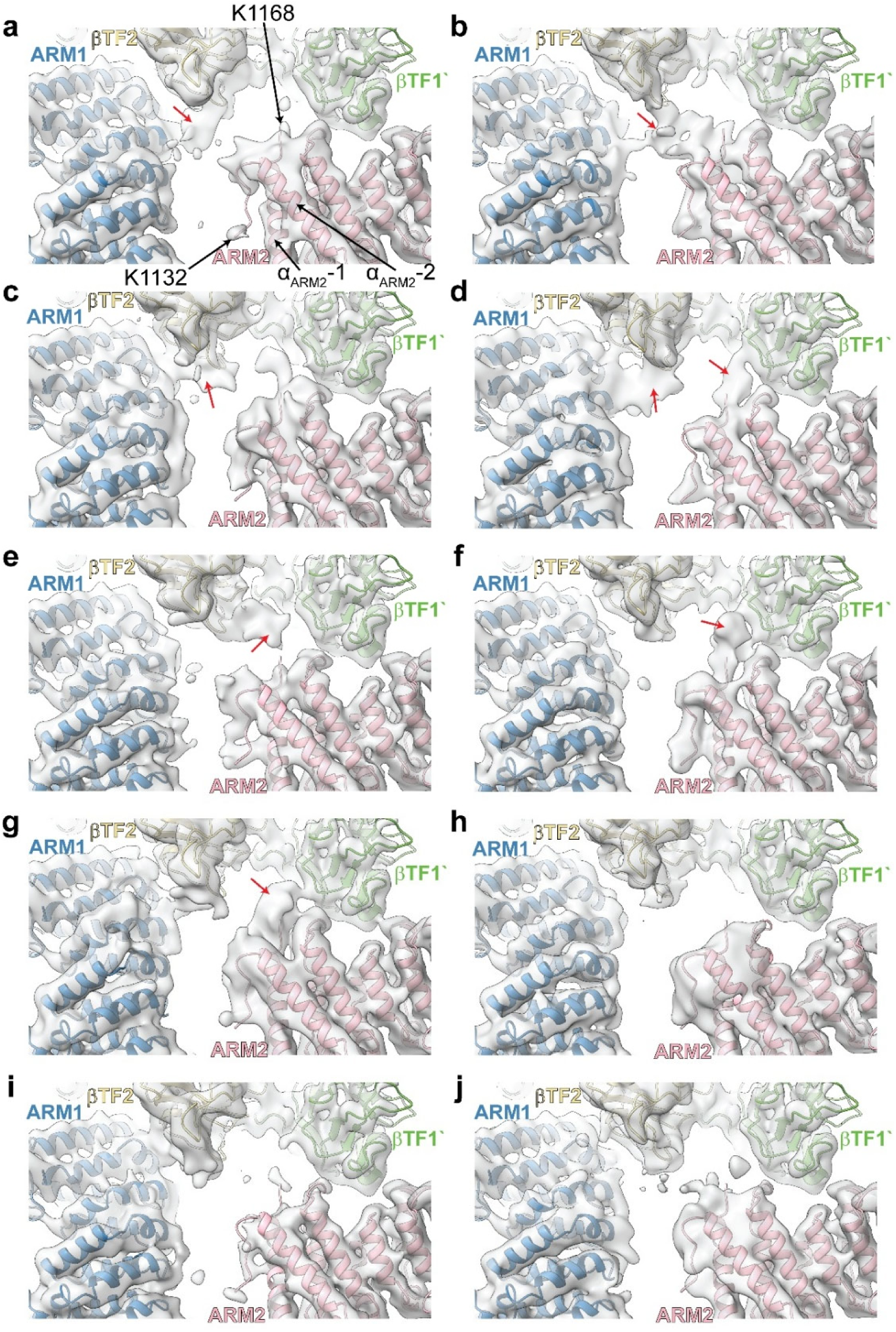
SBP dynamics in hIP_3_R-2. **a-j,** Focused 3D classification was performed with a mask encompassing the IP_3_-binding pocket after C4 symmetry expansion and signal subtraction. Representative class maps (low-pass filtered to 5 Å) are shown; red arrows highlight additional densities within the IP_3_ pocket or near the adjacent subunit. These features are consistent with alternative SBP configurations, including self-occlusion of the IP_3_ site and intersubunit engagement, although the maps were insufficient to model SBP residues unambiguously.

**Supplementary Fig. 4:**
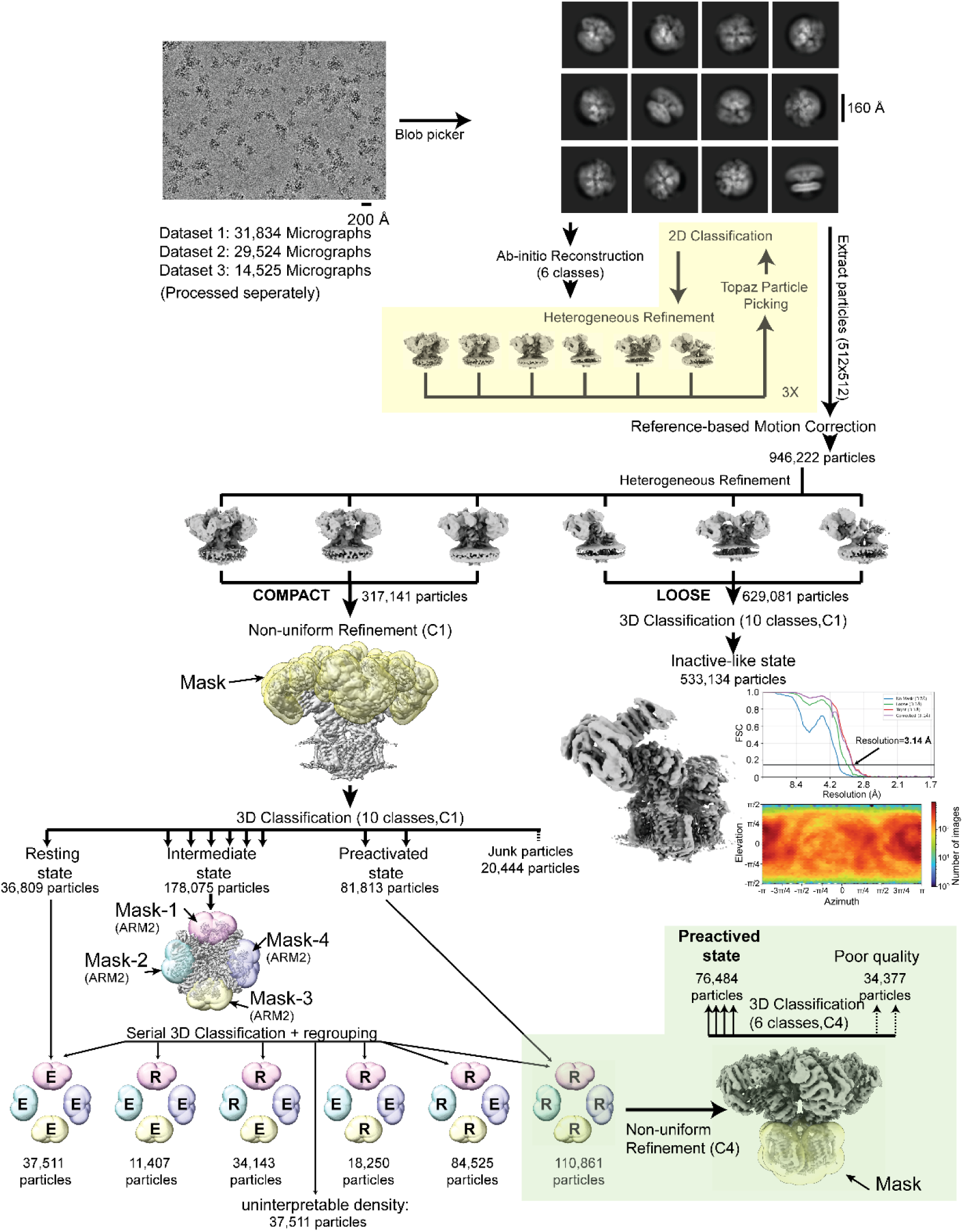
Cryo-EM analysis of hIP_3_R-2 in the presence of IP_3_ and ATP. Representative micrograph of hIP_3_R-2 supplemented with IP3 and ATP, reference-free 2D class averages, and the data-processing workflow from particle picking through 3D classification/ refinement, yielding compact and loose classes. Particles in the compact class were further subjected to sequential focused classification—first with a mask covering the entire cytosolic assembly, then with per-subunit masks on ARM2—yielding resting, intermediate, and preactivated classes. Validation for the inactive-like (loose) state includes gold-standard FSC curves (0.143 criterion) and particle angular distributions for the principal reconstructions. Validation plots for other classes are shown in Supplementary Fig. 5-7. E stands for extended ARM2, whereas R indicates retracted ARM2 conformations. See Methods for details on data acquisition and processing.

**Supplementary Fig. 5:**
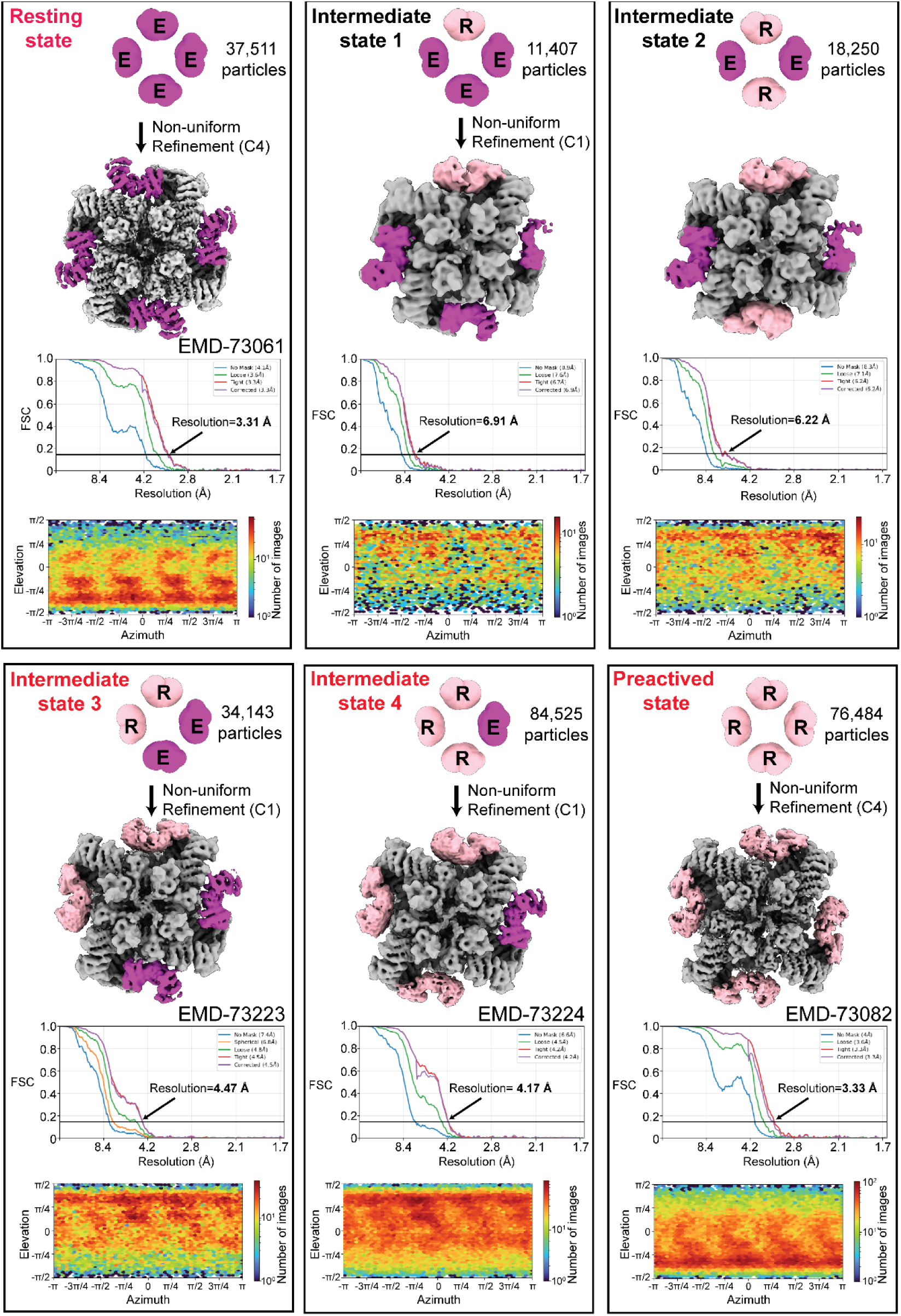
Structural heterogeneity of hIP_3_R-2 in the presence of IP_3_ and ATP. Cryo-EM reconstructions of the compact-class subclasses (resting, intermediate (1-4), and preactivated) are shown alongside gold-standard FSC curves (0.143 criterion) and particle angular distributions. ARM2 domains are colored to indicate conformation: extended (E; magenta) or retracted (R; light pink). EMDB accession codes are provided for deposited maps.

**Supplementary Fig. 6:**
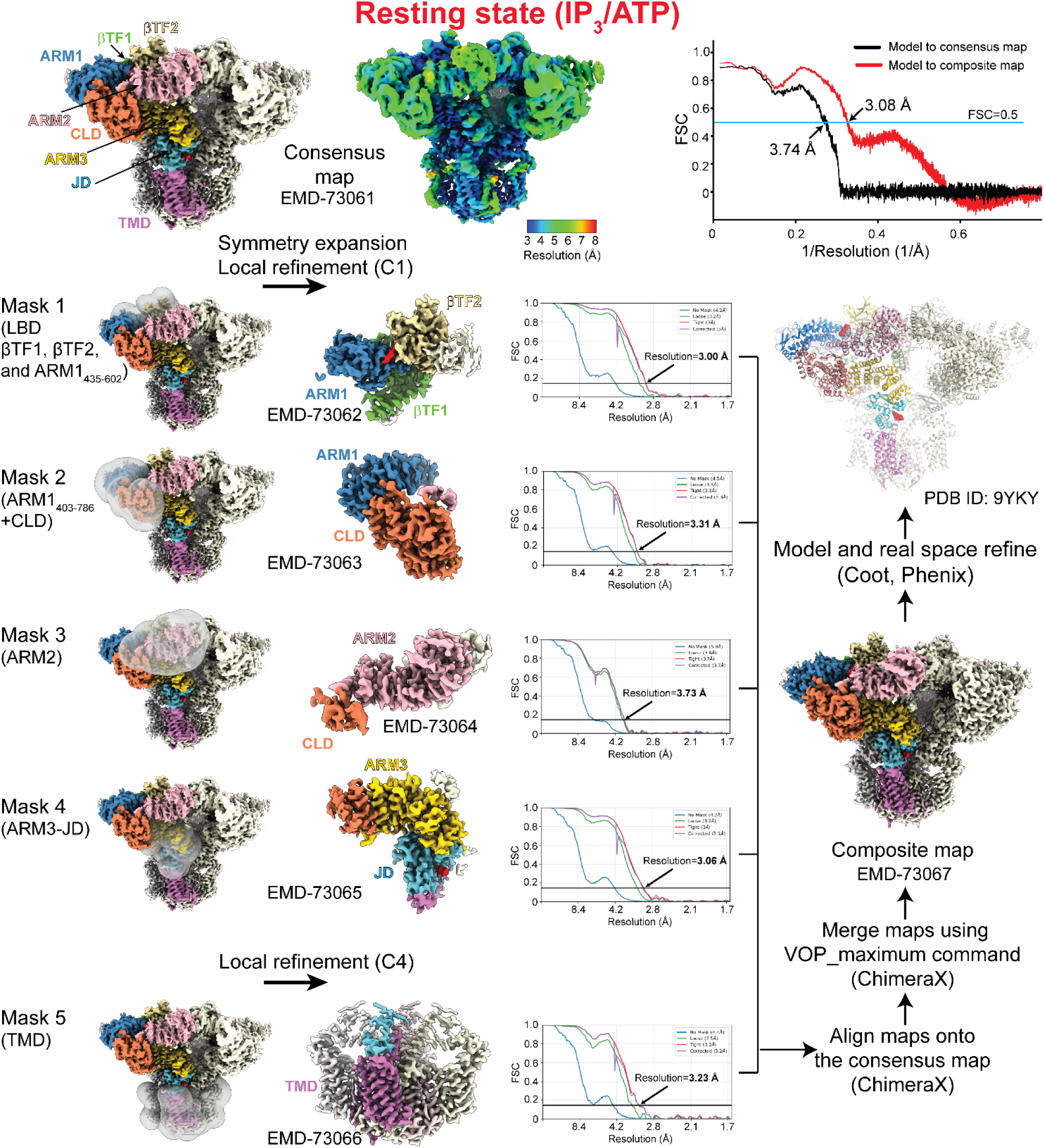
Local refinement of the hIP_3_R-2 in the resting state. Workflow from the consensus C4-symmetry reconstruction (3.31 Å overall) to symmetry expansion and focused refinements with per-subunit/per-domain masks, followed by assembly of a composite map from local refinements. Masks are shown as transparent overlays; domains are colored as in Fig. 1. Each locally refined map is labeled with its EMDB accession. For each mask, gold-standard FSC curves computed in cryoSPARC (0.143 criterion) are shown. Model-map FSCs compare the refined model against the consensus map (black) and the composite map (red).

**Supplementary Fig. 7:**
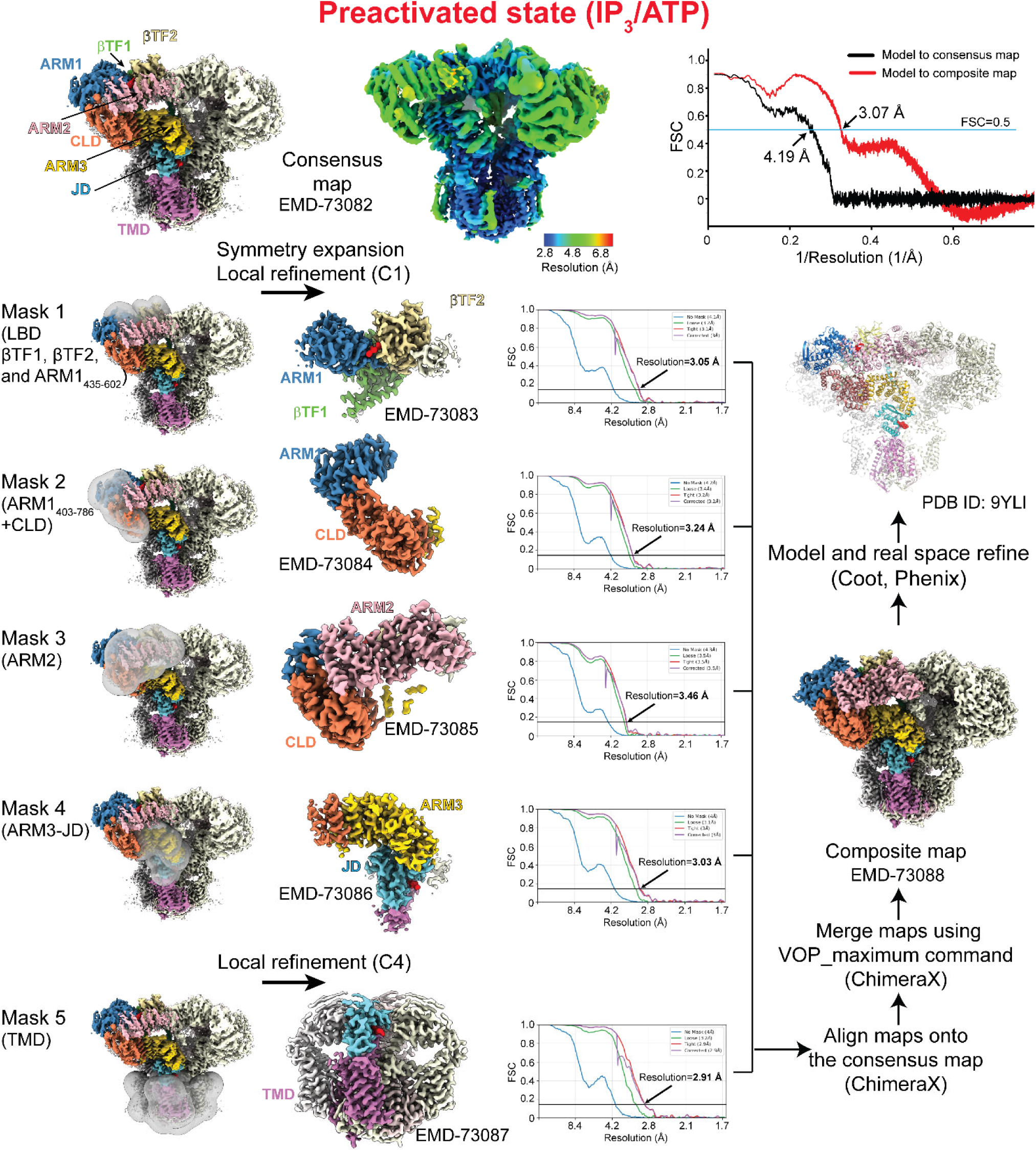
Local refinement of the hIP_3_R-2 in the preactivated state. Workflow from the consensus C4-symmetry reconstruction (3.33 Å overall) to symmetry expansion and focused refinements with per-subunit/per-domain masks, followed by assembly of a composite map from local refinements. Masks are shown as transparent overlays; domains are colored as in Fig. 1. Each locally refined map is labeled with its EMDB accession. For each mask, gold-standard FSC curves computed in cryoSPARC (0.143 criterion) are shown. Model-map FSCs compare the refined model against the consensus map (black) and the composite map (red).

**Supplementary Fig. 8:**
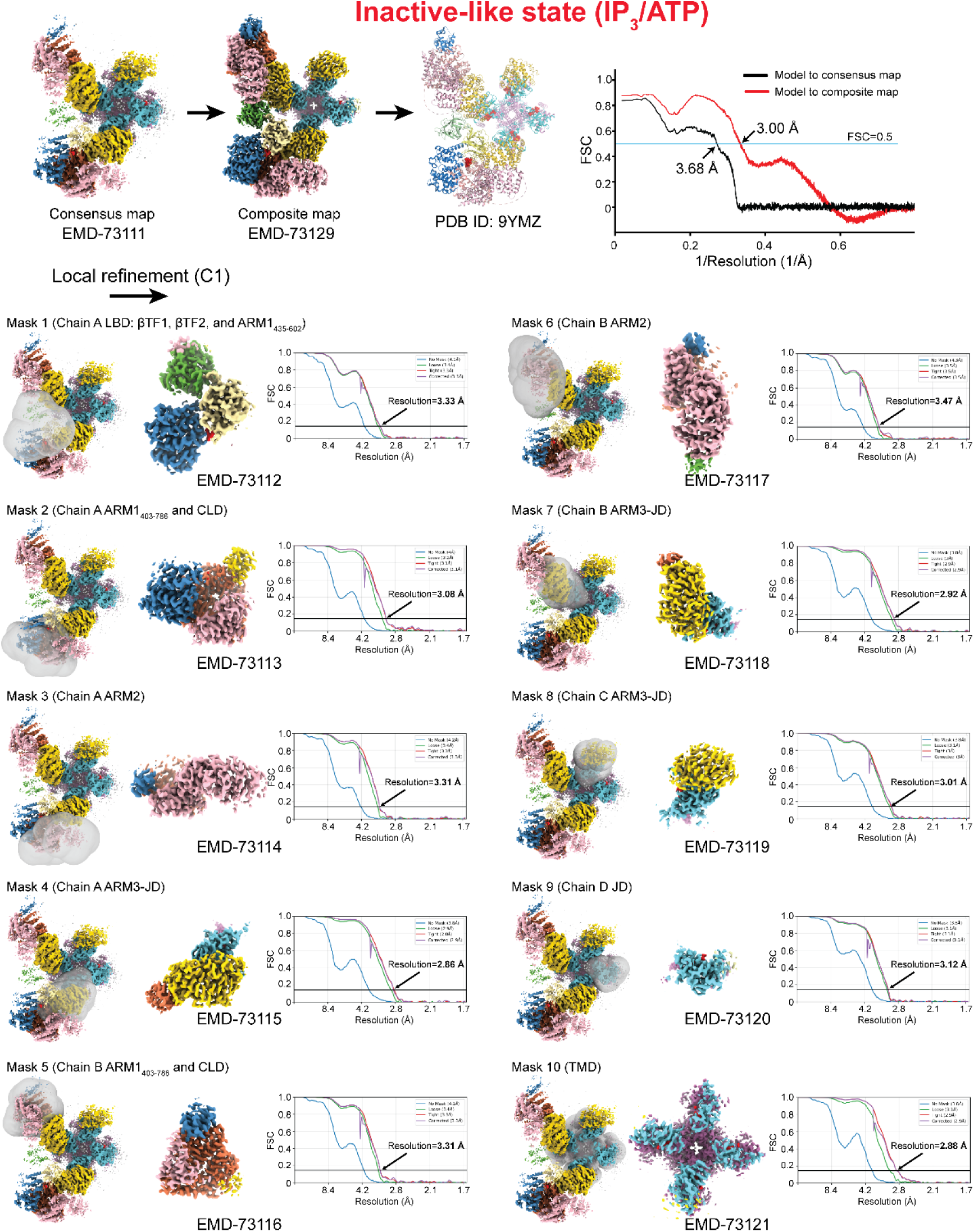
Local refinement of the hIP_3_R-2 in the inactive-like state (IP_3_/ATP). Workflow from the consensus reconstruction (3.14 Å overall) to focused refinements with per-domain masks, followed by assembly of a composite map from local refinements. Masks are shown as transparent overlays; domains are colored as in Fig. 1. Each locally refined map is labeled with its EMDB accession. For each mask, gold-standard FSC curves computed in cryoSPARC (0.143 criterion) are shown. Model-map FSCs compare the refined model against the consensus map (black) and the composite map (red).

**Supplementary Fig. 9:**
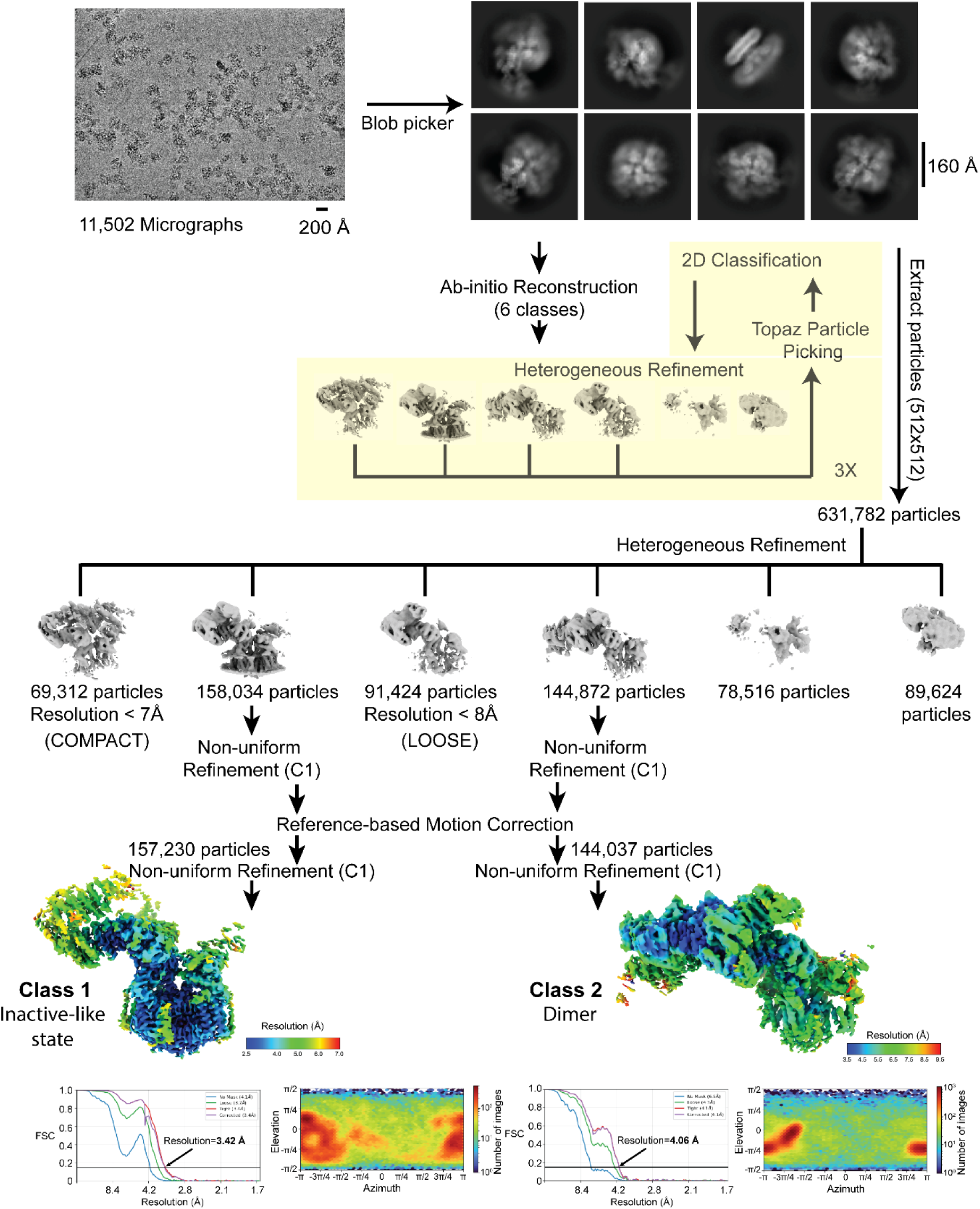
Cryo-EM analysis of hIP_3_R-2 in the presence of IP_3_, ATP, and Ca^2+^. Representative micrograph of hIP_3_R-2 supplemented with IP3, ATP, and Ca^2+^, reference-free 2D class averages, and the data-processing workflow from particle picking through 3D classification/ refinement, yielding two loose classes. Validation plots include local resolution maps, gold-standard FSC curves (0.143 criterion), and particle angular distributions for the principal reconstructions. See the Methods for details on data acquisition and processing.

**Supplementary Fig. 10:**
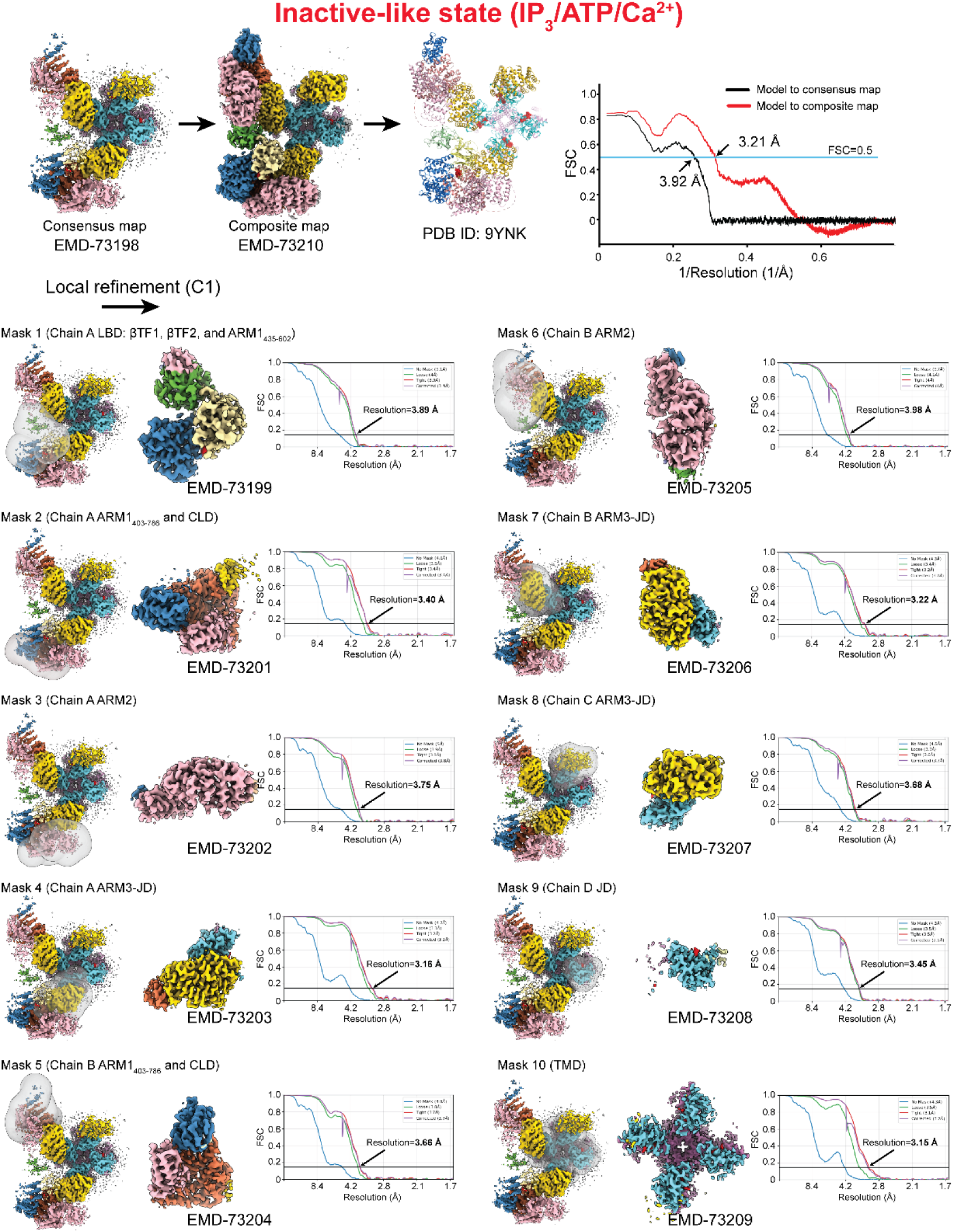
Local refinement of the hIP3R-2 in the inactive-like state (IP_3_/ATP/Ca^2+^). Workflow from the consensus reconstruction (3.42 Å overall) to focused refinements with per-domain masks, followed by assembly of a composite map from local refinements. Masks are shown as transparent overlays; domains are colored as in Fig. 1. Each locally refined map is labeled with its EMDB accession. For each mask, gold-standard FSC curves computed in cryoSPARC (0.143 criterion) are shown. Model-map FSCs compare the refined model against the consensus map (black) and the composite map (red).

**Supplementary Fig. 11:**
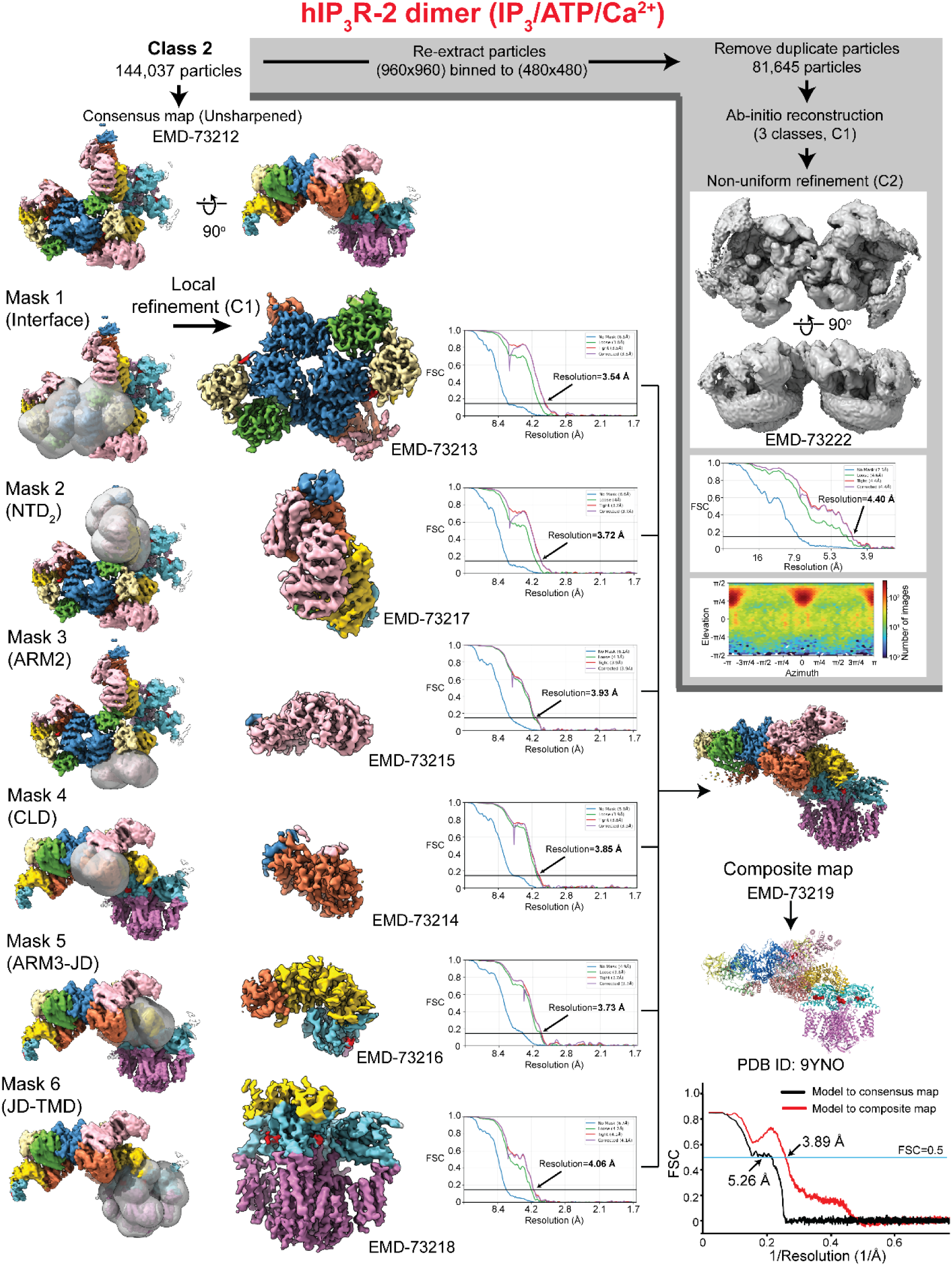
Processing of the hIP_3_R-2 dimer class. Two complementary workflows were applied to particles assigned to the dimer class. The first (gray-shaded panel) involves re-extraction with a larger box size to capture the full dimer, ab initio reconstruction, and 3D refinement to yield a global dimer map. The second proceeds from the consensus reconstruction (4.06 Å overall) through focused refinements with per-domain masks (including the interchannel interface) and assembly of a composite map from local refinements, improving local resolution at the interface to ∼3.5 Å and enabling modeling of contact residues. Masks are shown as transparent overlays; domains are colored as in Fig. 1. Each locally refined map is labeled with its EMDB accession. For each mask, gold-standard FSC curves computed in cryoSPARC (0.143 criterion) are shown, and model-map FSCs compare the refined model against the consensus map (black) and the composite map (red).

